# Conserved synteny reveals a saccharolytic lifestyle across *Kiritimatiellia*

**DOI:** 10.64898/2026.10.06.756941

**Authors:** Aaron Oliver, Sheila Podell, Eric E. Allen

## Abstract

*Kiritimatiellia* are a class of *Verrucomicrobiota* whose few cultured representatives are known for hosting large repertoires of carbohydrate-active enzymes and sulfatases. Many families of this class are restricted to metagenome-assembled genomes not yet discussed in literature, leaving their genome organization undescribed. Here, we survey 898 *Kiritimatiellia* genomes using similarity-colocalization networks (SCoNes), a data structure that captures conserved synteny among homologous genes across genomes. Using this approach, we investigate four metabolic adaptations to a glycan-rich environment: complex polysaccharide degradation, nitrogen cycling, hydrogenase activity, and natural product biosynthesis. Rather than unrelated functions, these capacities envelop a single anaerobic and saccharolytic strategy where sulfated glycan hydrolysis fuels fermentation, hydrogenases balance redox, and excess carbon supports nitrogen fixation.

**Importance:** Most microbes cannot be grown in the lab and are only identified by DNA fragments recovered from the environment, which makes it difficult to study the organization of their genomes. *Kiritimatiellia* are a class of sediment- and gut-associated bacteria that degrade and ferment complex sugars for energy, but our understanding of this lifestyle is based on a small number of cultured representatives. To understand how genes in core metabolic pathways are organized across hundreds of *Kiritimatiellia* genomes, we developed a new network analysis and visualization tool built to handle fragmented assemblies and flexible gene cluster boundaries. These network analyses reveal variable organization of catabolic enzymes and biosynthetic pathways, but a conserved core of metabolism that supports a lifestyle centered around polysaccharide utilization. These findings are of particular interest for understanding marine carbon cycling and discovering novel enzymes for the conversion of complex environmental sugars.

## Introduction

Annotation of well-characterized marker genes can reliably predict metabolic potential from metagenomes, but the organization of those genes remains difficult to study^1,2^. Detecting conserved genome organization requires methods that scale to hundreds of genomes, tolerate fragmented assemblies, and function independently of curated cluster boundaries or cultured relatives^3–5^. Here, we address these needs with similarity-colocalization networks (SCoNes), a new data structure for comparing gene organization across large sets of related genomes. We apply SCoNes to *Kiritimatiellia*, a class of mostly uncultivated *Verrucomicrobiota*, to understand how the organization of their genomes relates to biosynthetic and catabolic capacity.

This style of inference has a long history: genomic context has served as an early predictor of protein function for decades^6–8^, proving useful across problems as diverse as natural product discovery^9^, antibiotic resistance^10^, CRISPR-Cas systems^11^, methane consumption^12^, marine carbon cycling^13^, and the annotation of metagenomic dark matter^14^. Meanwhile, homology search tools have matured beyond basic pairwise BLAST searches into structure-aware^15^ and attention-based^16^ similarity metrics, each partitioning sequence space differently. The sequence similarity network (SSN) harnesses these distance metrics to visualize the sequence space of individual protein families^17^ or of entire genomes^18^. SSNs, through accessible webtools like EFI^19^ and EFI-GNT^20^, have been used to partition enzyme sets into functional subfamilies^21–23^ for small numbers of genomes, but this strategy struggles to parse sets of genes that work together in complex pathways or similar enzymes whose substrate specificity depends on neighboring auxiliary genes.

Network approaches already underlie homology and localization resources such as the STRING database^24^, but STRING encodes genomic context only for manually curated isolate genomes and relies heavily on prior experimental literature that most environmental lineages lack. Open-source tools that researchers can run on their own data fill more narrow niches: clinker^25,26^, antiSMASH^27^, and deepBGC^28^ are tailored to biosynthetic gene cluster (BGC) annotation and depend on predefined marker genes and gene-order constraints, making them poorly suited for whole-genome comparisons of syntenic regions. Other tools like FlaGs^29^ are more flexible in their metabolic scope but are rigid in their comparisons, interrogating a single query gene by considering only homologs from NCBI and immediate neighbors along a single contig. Despite this expanding toolkit, no single method visualizes conserved homologous gene neighborhoods across hundreds of genomes, a gap that becomes more consequential with the introduction of scalable whole-genome gene order comparison tools such as Spacedust^30^.

The SCoNe data structure developed in this paper is designed to avoid these limitations. A SCoNe begins as a sequence similarity network constructed using any pairwise homology metric, collapses gene nodes into communities of homologous genes, and links those communities based on genomic position. Because the gene neighborhoods are supplied as input rather than internally calculated, a SCoNe can be built using cluster boundaries from a flexible variety of sources; an annotation tool, synteny blocks inferred *de novo*, or the immediate neighbors of a single gene of interest. Comparing homologous sets rather than individual genes and evaluating colocalization across many genomes, is intended to keep the network interpretable at scale and tolerant of fragmented metagenomic assemblies. We illustrate each of these input types in the Results section, with case studies spanning different metabolic pathways associated with an anaerobic lifestyle.

In marine and gut-associated microbes, anaerobes are often linked to the breakdown of complex polysaccharides using colocalized collections of carbohydrate-active enzymes. Polysaccharide utilization loci are gene clusters marked by the presence of CAZymes^31^ and, in some environments^32^, sulfatase genes. Unlike BGCs, CAZyme gene clusters (CGCs) lack conserved gene order and consistent marker genes across phyla^33^, and they are disproportionately recovered from environmental samples whose fragmented, assembled contigs erode the performance of standard annotation tools^34^. Computational identification of CGCs is crucial for marine polysaccharides in particular, as fewer than 10% of experimentally validated CGCs act on marine substrates^35^ and many marine polysaccharide-degrading microbes remain uncultured. Leveraging this metagenomic dark matter requires scalable methods that can link uncultured lineages to their degradative potential.

One understudied yet significant degrader of marine polysaccharides are the *Kiritimatiellia*, a class of Verrucomicrobiota (or a potential new phylum^36^) with few cultured representatives. Although there are 23 taxonomic orders of sequenced *Kiritimatiellia* metagenome-assembled genomes (MAGs), all isolates come from the single order *Kiritimatiellales* found in marine and coastal sediments^37,38^. Publications about *Kiritimatiellales* center on their sulfatase-rich genomes, where sulfatases organized in CGCs^39^ can approach 10% of all coding sequences^40^. These CGCs are predicted to degrade sulfated algal polysaccharides such as agar, carrageenan, and fucoidan^41^. Some species have further been reported to fix nitrogen^38,42,43^ and detoxify antimicrobial compounds^40^ from marine polysaccharide producers. Whether these capacities represent independent adaptations or facets of one combined anaerobic strategy in *Kiritimatiellia* has not been explored. Here, we address this question by building SCoNes across nearly 900 public *Kiritimatiellia* genomes and benchmarking the effectiveness of SCoNes in visualizing and dividing enzyme families. Then, we investigate the anaerobic and saccharolytic lifestyle of this class through case studies in polysaccharide degradation, natural product biosynthesis, nitrogen fixation, and fermentative hydrogen metabolism.

## Methods

### Genome selection and annotation

All genomes of the class *Kiritimatiellia* were collected from the Genome Taxonomy Database version 226^44^, yielding 898 genomes and 2,203,257 predicted genes. Each genome was categorized into an environmental source (host-associated, aquatic & sediment, wastewater & built environment, or terrestrial soil) based on manual classification of NCBI BioSample metadata (**Supplemental Table 1**). Genomes were dereplicated to obtain a set of 480 species representatives^45^, but all predicted proteins from the *Kiritimatiellia* were leveraged for downstream analyses. A phylogenetic tree for all species representatives was constructed using GToTree v. 1.8.16, built upon MUSCLE v. 5.1 alignment and FastTree v. 2.1.11 phylogeny^46–48^, and visualized using the R v. 4.5.2 packages ggtree v. 4.0.5^49^ and ggtreeExtra v. 1.20.1^50^.

### Supplemental Table 1. Metadata and taxonomy of selected *Kiritimatiellia* genomes

Genomes were annotated for the presence of carbohydrate-active enzymes using the dbCAN database v. 11 with additional eCAMI annotations^51,52^. A subset of CAZyme annotations were classified into substrate-specific activity using the classification system proposed by López-Mondéjar et al.^53^. Sulfatase genes were annotated using a DIAMOND BLASTp^54^ screening using a coverage threshold of 0.35 and a minimum e-value of 1e-5 parameters, followed by the SulfAtlas v. 2.3.1 HMM webservice^39^ on initial hits.

Synteny blocks across all Kiritimatiellia were identified using Spacedust v. 2^30^. Nitrogen-fixation genes were annotated using pyHMMER^55,56^ from the Kofam^57^ and TIGRFAM^58^ HMM subset proposed by Pi et al.^59^. Hydrogenase genes were annotated using HydDB^22^. Sulfatase-aware CAZyme gene clusters, including relevant sugar transporters and regulatory genes, were identified using CGC-Finder^60^, while biosynthetic gene clusters were annotated using antiSMASH v. 8.0.4^61^ and BiG-SCAPE v. 2.0^9^ with c = 0.3.

## Network analyses

Sequence similarity networks were built in the visualization style of the EFI webtool^20^ using global DIAMOND BLASTp all-vs-all alignment of all 2,203,257 predicted proteins from the 898 Kiritimatiellia genomes. Graph manipulation and visualization was performed in Cytoscape.js^62^.

In the sequence similarity network, nodes represent individual proteins and two proteins are connected by an unweighted edge if they are within a pairwise similarity score e-value cutoff as determined by maximum closeness centrality^63^.

SSN communities of connected subgraphs (“homology communities”) were automatically detected using the Leiden algorithm^64^, and subgraphs of size 3 or less were dropped from the visualization. Homology communities were annotated using InterProScan v. 5.77^65^ on the most central member of each community as calculated by betweenness centrality using Scikit-network v. 0.33.5^66^ for interpretation of protein function. This includes the annotation of transporters and regulatory enzymes discussed in Case Studies 3 and 4.

## SCoNe construction

The process of generating a simplified similarity-colocalization network from the underlying SSN and gene neighborhood definitions is illustrated schematically in **Figure 1**. In this example, the initial sequence similarity network (G_SSN_) (**Figure 1b**) is generated using a pairwise BLASTp search, with e-value as the homology metric and alignment locations as a source of gene coordinates defining relative gene position (**Figure 1a**). A community detection algorithm divides the sequence space into communities based on pairwise distances (G_community_) (**Figure 1c**). Then, the colocalization step generates an additional edge type between genes that colocalize in user-defined neighborhoods (G_neighborhood_) (**Figure 1d**). Finally, homology communities are collapsed into nodes and colocalization edges between communities combined to form the edges of GScoNe (**Figure 1e**). A SCoNe construction script that accepts a list of pairwise homology scores and gene colocalization edges and outputs a Cytoscape^67^ visualization file is provided at https://github.com/AaronAOliver/KiritimatiallesSCoNe.

**Figure 1.**
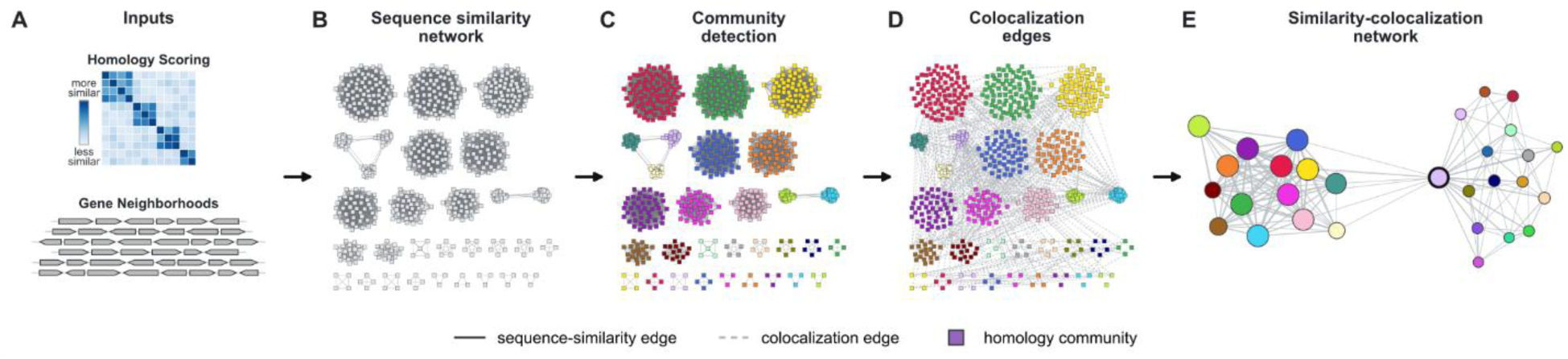
The multistep construction of a similarity-colocalization network using mock data. (A) Pairwise homology scores and gene coordinate data are sufficient to construct a SCoNe. (B) A sequence similarity network built from pairwise homology scores. (C) Clustering protein space through community detection. (D) Appending colocalization edges from gene cluster boundaries or anchored genome neighborhoods. (E) Collapsing homology communities into nodes to generate a similarity-colocalization network. The central pink node with two distinct subgraphs showcases unique gene contexts and potentially promiscuous biochemical function.

## Results

### Polysaccharide degradation repertoires in metagenomic *Kiritimatiellia*

To demonstrate the utility of SCoNes on metagenomic datasets, where contigs are often short and gene clusters fragmented, we collected all *Kiritimatiellia* genomes from the Genome Taxonomy Database and surveyed their capacity to degrade polysaccharides. Metagenome-assembled genomes of the *Kiritimatiellia* derive primarily from environmental and gut microbiomes, and only a handful of strains have been cultured to date^38^. The phylogenetic distribution of species representatives and their carbohydrate-degradation capacity is shown in **Figure 2**, with counts for a subset of CAZyme families of validated substrate specificity^53^ plotted beneath the tree. Two families stand out for their exceptional sulfatase and algal-polysaccharide-degrading capacity and are highlighted on the tree: marine *Pontiellaceae* and freshwater CAIQIC01.

**Figure 2.**
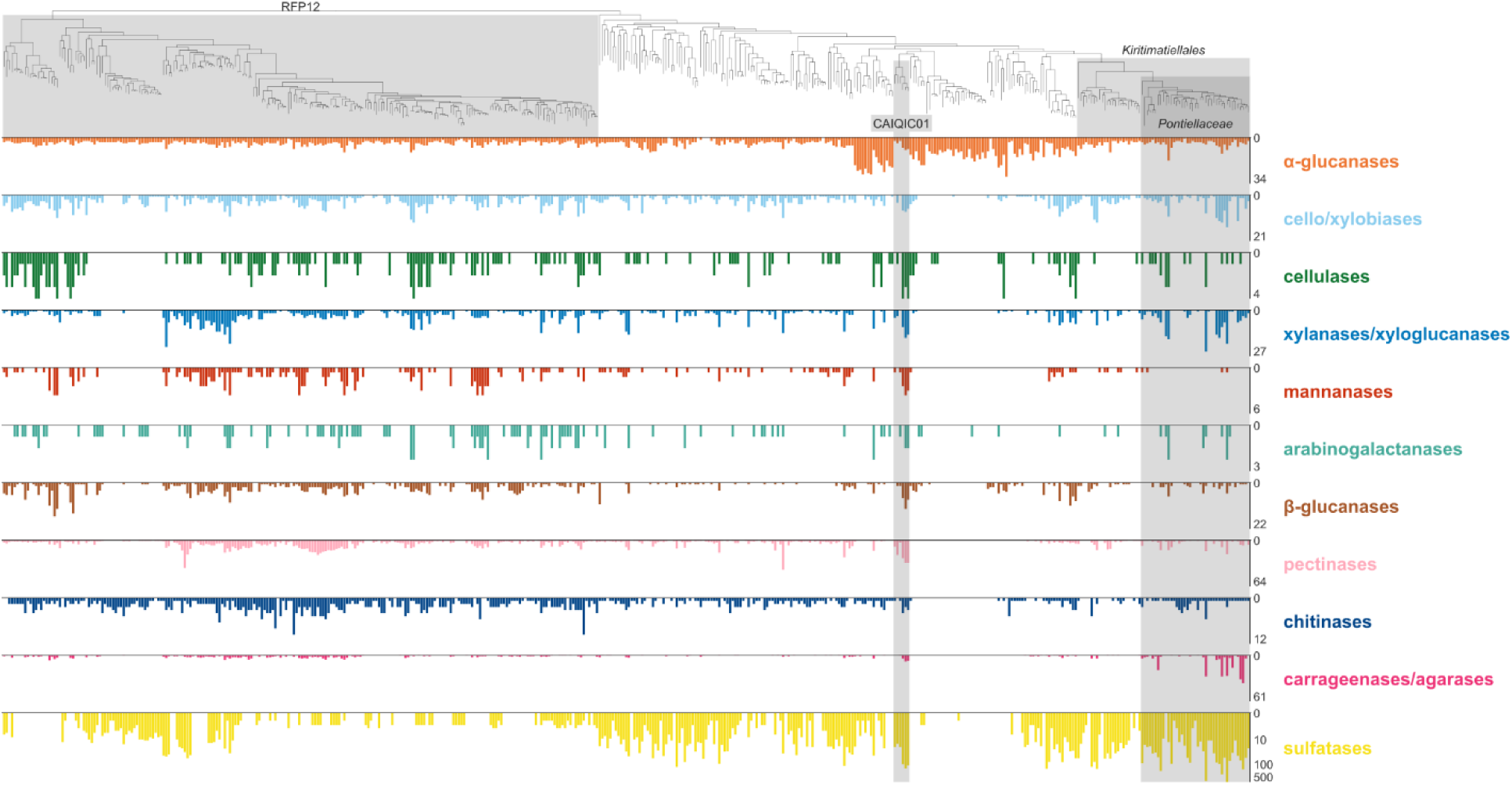
Polysaccharide degradation potential of the *Kiritimatiellia*. On top, a multigene marker phylogenetic tree of 480 species representative *Kiritimatiellia* genomes from the Genome Taxonomy Database. Two selected families with diverse enzyme repertoires, *Pontiellaceae* and CAIQIC01, highlighted in gray. Subsequent rings denote counts of catalytic carbohydrate enzymes and sulfatases per genome. Due to expanded sulfatase counts for some genomes, the bottommost ring uses a log scale.

Carbohydrate utilization potential aligns closely with environmental source in our dataset (**Supplemental Figure 1a**). Aquatic genomes are enriched in xylanases and sulfatases, while host-associated genomes have more cellulases, chitinases, and glucanases, consistent with environmental polysaccharide distributions. Marine *Pontiella* are outliers among aquatic *Kiritimatiellia*, with specialization toward sulfated algal polysaccharides such as carrageenan and agar (**Supplemental Figure 1b**). Individual genomes from *Pontiellaceae* have previously been found on macroalgal surfaces^43,68^. The cultured *P. desulfatans* and *P. sulfatireligans* encode 500 and 461 sulfatase genes respectively, more than any metagenome-assembled genome in the dataset. The freshwater CAIQIC01 are generalists, with a median of 217 CAZymes and 101 sulfatases per genome, the highest median sulfatase density of any family. Primarily found in lake and aquifer metagenomes^69^, uncultured CAIQIC01 combine xylan- and sulfated-polysaccharide degradation repertoires with larger amounts of cellulases, glucanases, and other enzymes acting on labile sugars.

## Case study 1: polysaccharide utilization gene clusters

To explore the genomic organization of polysaccharide degradation strategies of the *Kiritimatiellia*, CAZyme gene clusters (CGCs) were annotated in our genome set using dbCAN. In total, 2,298 gene clusters were identified, containing 4,825 CAZyme genes, 3,443 sulfatases, 4,522 transporters, 3,383 auxiliary genes, and 6,811 hypothetical genes. Multiple CAZymes were found in 44.0% of CGCs, with at least one sulfatase in 63.3%, and multiple sulfatases in 26.8%. We used the gene cluster boundaries as the basis of colocalization edges, and DIAMOND e-values for pairwise homology scores for all genes within a CGC. We then asked whether integrating genomic neighborhood information could identify conserved gene families whose organization is underexplored relative to well-annotated carbohydrate-active enzymes.

First, an e-value cutoff threshold of 1e-80 was chosen to construct a sequence similarity network of several thousand proteins (**Supplemental Figure 2**). Collapsing this sequence-similarity network into homology communities yielded the SCoNe in **Figure 3a**, in which each CAZyme, sulfatase, transporter, and regulatory family collapses to a small number of communities whose genomic neighborhoods can be examined individually. In total, the full set of CGC genes could be condensed into 188 homology communities of size 5 GCGs or larger. Sulfatases and auxiliary genes showed wider variance in sequence diversity and genomic context than CAZyme families.

**Figure 3.**
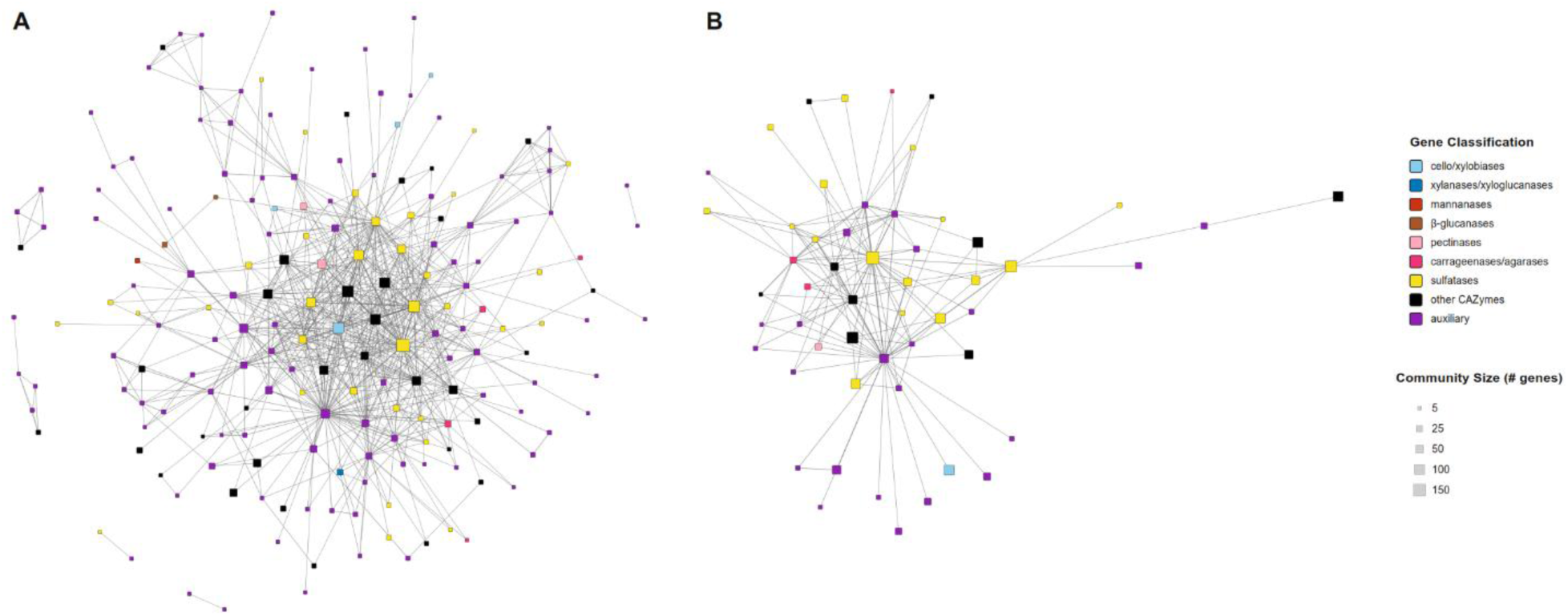
Similarity-colocalization network of conserved genes in CAZyme gene clusters of *Kiritimatiellia*. (A) The entire SCoNe built from dbCAN-annotated CGCs. (B) The SCoNe pruned to focus on genes associated with predicted agarases and carrageenases. Edges that colocalize in at least 5 CGCs shown. Singleton nodes and unannotated subgraphs are not shown.

Although a full SCoNe can span thousands of genes, the interpretability of the entire network at once is often quite difficult. One benefit of SCoNes is that network subgraphs or a selection of just a few nodes can be extracted, highlighting more detailed relationships for later ad-hoc exploration. Pictured in **Figure 3b** is a subset of the SCoNe in **Figure 3a**, looking only at genes colocalized with a carrageenase or agarase. Nodes in the center of each network colocalize with the highest diversity of genes, denoting core or multifunctional enzymes or auxiliary genes. This pruned SCoNe, containing only 42 homology communities, resolved a catalytic core of glycoside hydrolase subfamilies (including GH86, GH167, GH50, GH82, and GH117) frequently colocalized with more than a dozen sulfatase families (including S1_7, S1_15, S1_16, S1_17, S1_19, and S1_20) and 23 auxiliary families, whose conserved colocalization to the catalytic CGC core had not previously been appreciated. These conserved auxiliary families include primary (ATP-binding cassette) and secondary (major facilitator superfamily, solute:sodium symporter) sugar transporter families, two-component regulator systems previously shown to regulate polysaccharide utilization loci in marine bacteria^31,70^, and formylglycine-generating enzymes required to post-translationally activate S1 sulfatase enzymes^71^. Both SCoNes pictured in **Figure 3** are available as Cytoscape visualization files at Zenodo 10.5281/zenodo.22103458.

## Case study 2: aryl polyene biosynthetic gene clusters

Biosynthetic gene clusters also serve as a relevant use case for visualizing hundreds of divergent gene neighborhoods at scale. Aryl polyene clusters were the second most abundant biosynthetic class across Kiritimatiellia species representatives, comprising 14.2% of total BGC annotations, exceeded only by terpene and terpene-precursor clusters (**Supplemental Figure 3**). The SCoNe pipeline condensed these genomic neighborhoods into 43 conserved SSN communities present in at least 10 clusters (**Figure 4a**). The network’s central members consisted of two β-ketoacyl synthases and a ketoreductase family, with separate clusters within the SCoNe containing additional biosynthetic domains unique to separate gene cluster families. Two of the larger aryl polyene gene cluster families illustrate how auxiliary enzymes organize around a central backbone: GCF-3 (**Figure 4b**) and GCF-2 (**Figure 4c**). These two GCFs share a core set of five genes (two ketosynthases, a ketoreductase, an acyltransferase, and a dehydratase), where GCF-2 adds a hydrolase gene and GCF-3 includes a more complex cassette of two acyl carrier proteins, a lipoprotein carrier, and an MmpL-like transporter. Both GCFs were primarily found in UBA1067, an uncharacterized genus linked to human^72^ and rumen^73^ gut microbiomes.

**Figure 4.**
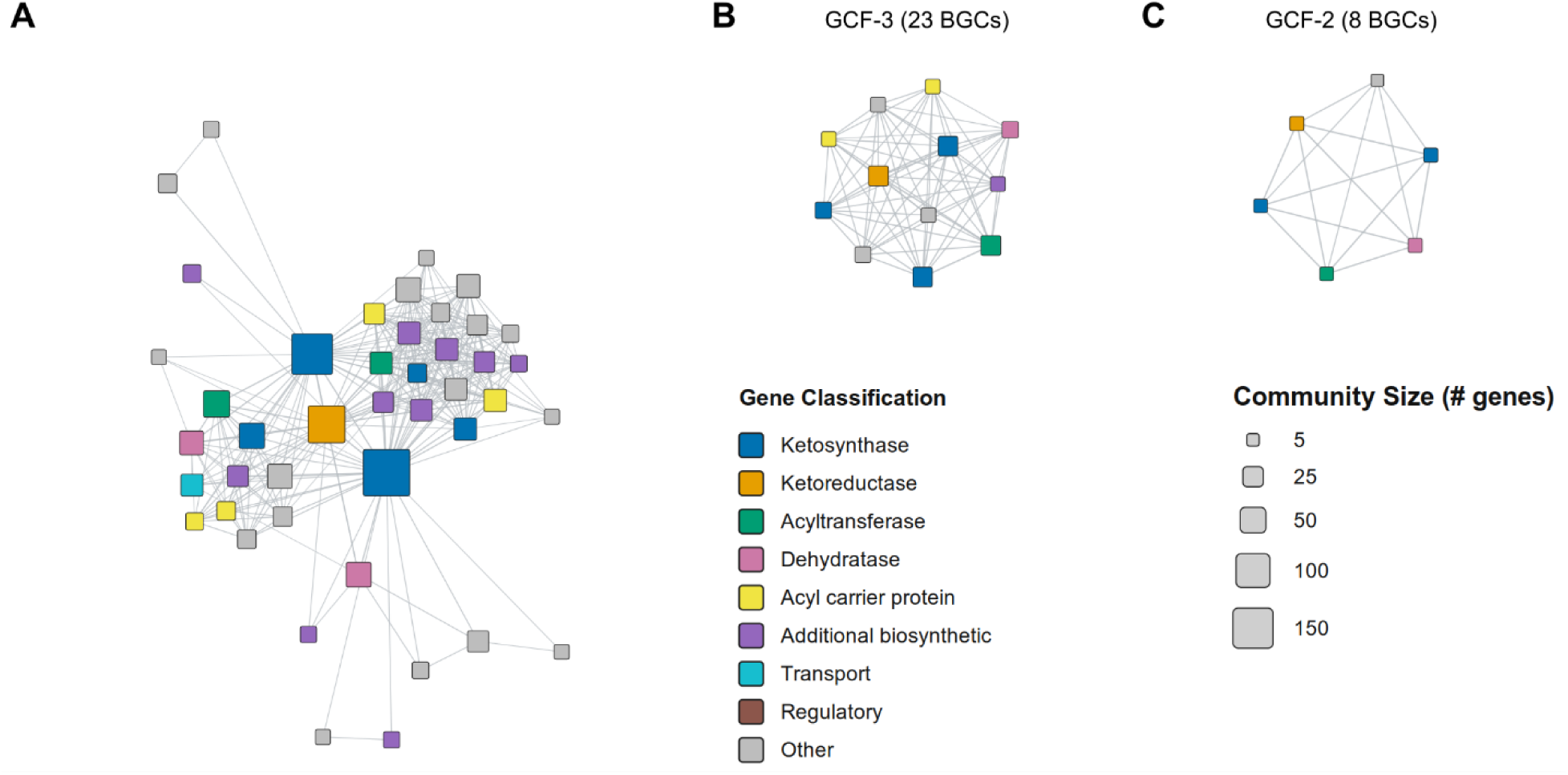
Biosynthetic potential of *Kiritimatiellia*. (A) A SCoNe of conserved aryl polyene biosynthesis genes shows a consistent backbone. Nodes are colored by antiSMASH annotation class and sized based on the number of homologous genes. (B, C) Pruning the main SCoNe produces SCoNes of individual gene cluster families.

## Discovery and analysis of conserved synteny blocks

While visualization strategies that scale to hundreds or thousands of genomes are particularly useful for gene clusters with established annotation tools, such as biosynthetic or CAZyme gene clusters, the boundaries of other gene neighborhoods are less standardized.

Recent tools involving gene orthology and neighborhood now allow the discovery of conserved gene synteny strategies without requiring specialized databases or a priori gene-cluster boundaries. One example of these tools is Spacedust^30^, which uses Foldseek and MMseqs2 to identify conserved blocks of orthologous, colocalized genes across many genomes. **Supplemental Figure 3** applies this to the 6,505 genes of the cultured *Pontiella desulfatans* genome^37^ scored for conserved synteny across 479 species-representative *Kiritimatiellia* genomes, 41 of which are within the family Pontiellaceae (**Supplemental Figure 4a**) and 438 across the rest distributed across the order (**Supplemental Figure 4b**). Few gene neighborhoods are conserved across the entire order (**Supplemental Figure 4c**). Instead, the conserved blocks partition those retained within *Pontiella* and those shared more broadly, revealing lineage-specific genome organization. Because these blocks require neither pre-defined neighborhood boundaries nor core marker genes, the approach scales to the organization strategies of hundreds of related genomes at once.

## Case study 3: nitrogen fixation operons

One region of homologous genes highlighted by synteny block analysis is the nitrogen fixation (nif) operon. *Kiritimatiellia* have been shown to fix nitrogen in culture^38^, and reference-free synteny analysis identified a nif gene cluster in 31 genomes across the class, 20 of which carry the complete nitrogen fixation catalytic core (**Figure 5**). Four marine Pontiella genomes, *P. sulfatireligans*^37^ and three uncultivated representatives, encode a complete molybdate ABC transporter colocalized with the larger operon of Mo-dependent nitrogenases (**Figure 5c**). Despite the contig fragmentation common in metagenomic sequencing, the SCoNe approach tolerates unconserved gene order and still captures the operons with variable gene content and missing marker genes (**Figure 5b**). For example, several *Pontiella* genomes (e.g., *P. agarivorans*) carry *nifENB* upstream rather than downstream of *nifHDK*, and metagenome-assembled genomes frequently have molybdate transport in distinct operons from the primary nif cluster.

**Figure 5.**
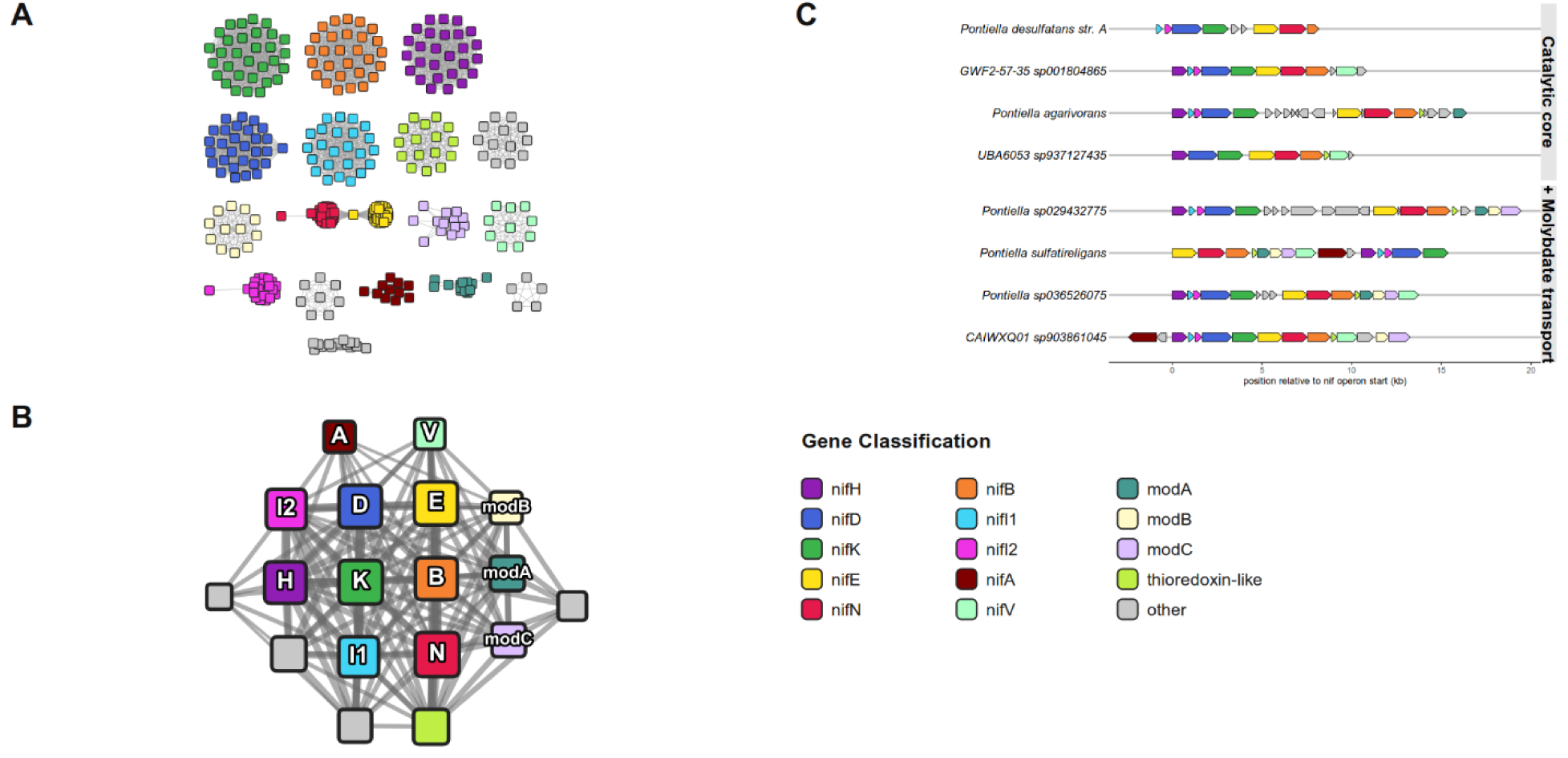
Paired SSN and SCoNe of nitrogen fixation gene clusters in the *Kiritimatiellia*. (A) A sequence similarity network of *nif* gene operons. (B) A SCoNe showing organization of *nif* and molybdate transport clusters, using the same color scheme as (A). (C) A gene neighborhood map showing two styles of *nif* operon synteny.

## Case study 4: [FeFe]-hydrogenase loci and fermentative hydrogen metabolism

Synteny block analysis also identified a [FeFe]-hydrogenase locus conserved across Kiritimatiellia (**Figure 6**), typically a trimeric *hydABC* operon accompanied by a small [2Fe-2S] ferredoxin. 78% of the Kiritimatiellia genomes analyzed encode at least one [FeFe]-hydrogenase, and many encode more than one type: the electron-bifurcating Group A3 enzyme is abundant (74% of all genomes), while the sensory *hydS*-containing Group C (32%), ancestral Group B (30%), and fermentative Group A1 (5%) types occur less frequently. A SCoNe built from all hydrogenase loci (**Figure 6b**) resolves this diversity into distinct operon architectures. In the *hydS* architecture, *hydB* is absent and the sensory *hydS* colocalizes with a PP2C-family phosphatase and an alcohol dehydrogenase, consistent with signal transduction and control of fermentative pathways. Elsewhere, the *hydB* subunit is instead extended: in 163 genomes it fuses a FAD/NAD(P)-binding oxidoreductase module and additional [4Fe-4S] clusters onto the ancestral core, and 132 genomes carry both the fused and the unfused form. The cultured *K. glycovorans* and *T. aerotolerans* genomes carry the fused *hydB* within a tandemly duplicated *hydABC* array, adjacent to but distinct from its canonical unfused operon (**Figure 6a**).

**Figure 6.**
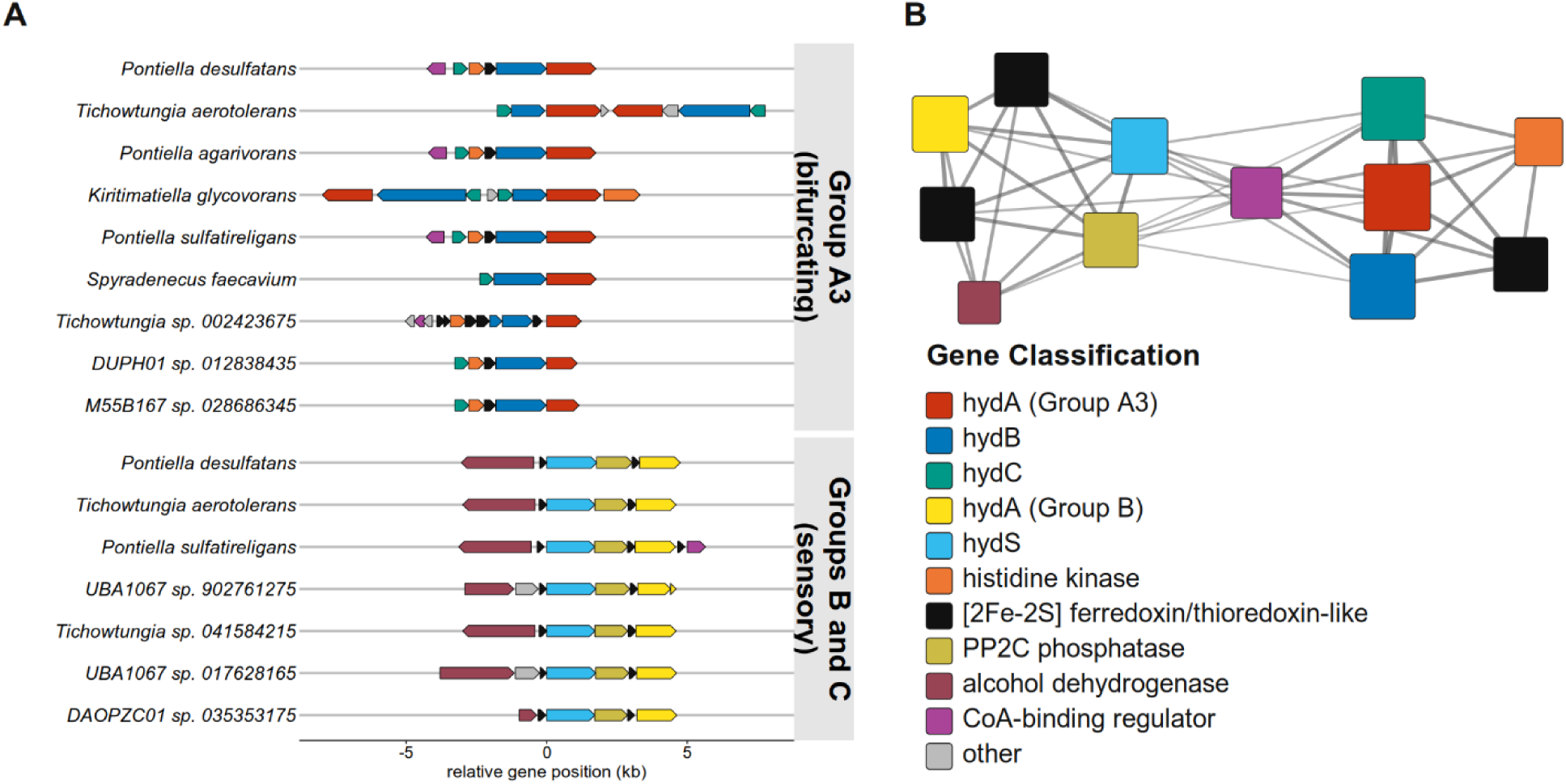
Genomic organization of [FeFe]-hydrogenase synteny blocks. (A) Gene neighborhood diagram of selected genomic regions containing the hydrogenase cassette *hydABC* or standalone hydrogenase *hydS* and conserved auxiliary genes. (B) A similarity-colocalization network of genes associated with [FeFe]-hydrogenase, following the same color scheme as (A).

## Discussion

In this study, we present the first class-wide investigation of the potential of Kiritimatiellia to degrade terrestrial and marine polysaccharides. We have identified trends across the phylogenetic sub-groups of the Kiritimatiellia, highlighting families with unique metabolic potential to degrade a wide variety of glycan substrates. We also confirm that several biochemical processes previously limited to a few cultured isolates, such as nitrogen fixation, hydrogen production, and natural product biosynthesis, are widespread among uncultured *Kiritimatiellia* genomes.

For glycan degradation, we see clear differentiation in enzyme repertoire by environmental source. In particular, sulfatase enzymes are enriched in genomes belonging to bacteria that live in aquatic environments and marine sediment. This is consistent with the enrichment of sulfated monosaccharides in algal sugars commonly found in the ocean. Sinking marine debris also explains the enrichment of these enzymes in sediment microbes as it is well established that sediment microbes play an important role in cycling carbon of particulate algal biomass^42^. While most *Kiritimatiellia* in our dataset are rich in CAZymes, two groups stand out in their adaptation towards degrading algal polysaccharides, as marked by an abundance of sulfatase enzymes.

The first, the family *Pontiellaceae*, is the most well-described lineage across all of the *Kiritimatiellia* and includes several cultured isolates. *Pontiella* have been isolated from anoxic marine sediments and microbial mats^38,74^, and our analysis of uncultured microbes shows that this algal polysaccharide targeting is consistent across the family regardless of environmental source. Still, the isolate *Pontiella* genomes do contain a very high number of sulfatase enzymes, higher than any *Pontiella* MAG (**Figure 2**). **Supplemental Figure 1b** further displays the unique repertoire of carbohydrate degradation genes of Pontiella relative to the rest of the class, driven by this expansion of sulfatases together with an abundance of carrageenases and agarases. This specialization is further apparent as the relative abundance of glucanases and chitinases in Pontiella is far lower than in Kiritimatiellia from wastewater and mammalian gut microbiomes.

On the other hand, the undescribed metagenomic CAIQIC01 family is composed solely of freshwater microbes from water column or freshwater sediment samples^69^ that are simultaneously rich in sulfatases and in xylanases, pectinases, chitinases, and cellulases. This sulfatase abundance also extends to some host-associated lineages, which does serve an important biological purpose, as host mucins and glycosaminoglycans are often heavily sulfated^32^. For example, UBA1067, found in mammalian gut microbiomes^73,75^, contains up to 11 sulfatases per genome, and two UBA6053 genomes recovered from coral mucus^76^ contain 80 and 82 sulfatases. Members of these uncultured families carry sulfatase levels otherwise seen only in *Pontiella* and a few Verrucomicrobiota^13,77^, yet they have never been described in prior literature as a potential source of novel polysaccharide degradation strategies.

*Pontiella* also display the ability to fix nitrogen using a molybdenum-dependent system. The relationship between sulfated polysaccharide degradation and nitrogen fixation may stem from large amounts of carbon brought on by an influx of sinking particulate organic matter, raising local C:N ratios^78^. The bottleneck of nitrogen supply can be bypassed using the excess carbon and energy provided by this set of recalcitrant polysaccharides. Other heterotrophic marine bacteria, including Verrucomicrobiota and Proteobacteria, likewise raise their rates of nitrogen fixation in the presence of bioavailable mono- and polysaccharides^79–81^. In the eastern Mediterranean, addition of glucose increased nitrogen fixation by up to tenfold^82^, an effect attributed to the high C:N, low-oxygen microenvironments created by carbon-rich marine particles. Carbon is also readily available in the phycosphere due to glycan seepage by kelp^83^, and one study found that *Pontiella* represented up to 45% of nifH gene activity of the kelp-surface microbiome, following a diurnal cycle^43^. Previous studies have shown that macroalgal blooms select for heterotrophic nitrogen fixers as macroalgae utilize nitrogen in the water column^84^ but provide a niche for polysaccharide degraders.

Complex polysaccharide degradation is only one component of an anaerobic lifestyle, and our survey shows strong genomic evidence of fermentation across Kiritimatiellia. [FeFe]-hydrogenases appeared in most species representative genomes, with conserved auxiliary genes immediately surrounding the hydABC cluster. Previous analysis of bifurcating hydrogenases noted that *P. sulfatireligans* and *P. desulfatans* appeared to lose the hydD subunit present in related phyla such as *Planctomycetota* and *Hydrogenedentota*^85^. This analysis extends that observation, as hydD is absent across the class, supporting the hypothesis that hydD loss preceded diversification of the class rather than occurring independently in Pontiella. However, tandem duplication of the fused hydB subunit in *K. glycovorans* and *T. aerotolerans* suggests hydrogenase metabolism continues to diversify in this lineage^86^.

Under the metabolic model described by our case studies, sulfatases and glycoside hydrolases release monosaccharides from algal and host glycans, fermentation extracts energy from those monosaccharides, and hydrogenases discharge those electrons as H_2_. This coupling is explicit in *P. agarivorans,* where approximately 75% of fermentation electrons are recovered in organic products and the remainder are available for nitrogen fixation, with hydrogen gas appearing as a minor fermentation product^38^. Aryl polyenes, known to quench reactive oxygen species and support biofilm formation^87,88^, shield these anoxic processes from the outside environment.

Enzymes acting on sulfated polysaccharides are of considerable interest for the valorization of algal biomass^89^, and targeted culturing of novel *Kiritimatiellia* for enzyme discovery has been established by our group in bioreactor systems^40^. Uncultured marine microbiomes are similarly a continuing source of novel natural products^90^. Our survey represents the first comprehensive study of polysaccharide utilization and natural product potential across Kiritimatiellia, which should serve as a useful reference when further mining these genomes for enzymes relevant to bioenergy and new pharmaceuticals. The gene neighborhood approach enabled by SCoNe in this manuscript is particularly effective in highlighting conserved hypothetical and novel genes in widespread gene clusters like CGCs and BGCs, which would otherwise require manual scanning to find putatively novel enzymes^91^.

We have demonstrated that SCoNes are useful at three levels of gene neighborhood characterization. First (**Figure 3**), SCoNes apply to well-defined gene cluster boundaries produced by established tools. This requires only the gene sequences within each cluster and their relative arrangement, so transforming the output of tools such as dbCAN, antiSMASH, or deepBGC into a SCoNe demands little additional computation and can visualize the metabolic or catabolic potential of hundreds or thousands of genomes at once, which popular gene arrangement plots like clinker^25^ cannot match at scale. Second (**Figure 5**), because SCoNes do not require predetermined cluster or neighborhood boundaries, they are especially suited to the conserved gene neighborhoods of biochemically important regions, regardless of the existence of specialized annotation tools. Basic gene organization and similarity metrics quantified with pairwise homology tools are sufficient to identify and analyze synteny blocks in a scalable way. Third, a SCoNe visualization can quickly predict multifaceted activity of a single gene based on multiple conserved gene neighborhoods. In our hydrogenase vignette (**Figure 6**), the annotation of a single *hydA* gene was sufficient hypothesis generation to subset the Kiritimatiellia SCoNe and identify two divergent gene cassettes for bifurcating and sensory hydrogenases.

## Conclusion

These analyses confirm *Kiritimatiellia* as anaerobic and saccharolytic specialists with metabolism organized around breaking down sulfated glycans. These bacteria live in low-oxygen environments, such as marine sediments or animal guts, and interface with complex, often sulfated polysaccharides. In response, *Kiritimatiellia* hydrolyze large glycans into simple monomers for fermentation, using hydrogenases to balance redox. Some lineages, particularly *Pontiella*, divert energy from fermentation into molybdenum-dependent nitrogen fixation when bioavailable nitrogen is scarce. Describing the breadth of glycan cycling strategies in uncultured marine lineages is crucial for understanding the fate of carbon in the oceans.

The novel SCoNe data structure offers a scalable way to visualize and explore genome organization directly from environmental assemblies. This approach succeeds when cluster boundaries are defined by specialized tools, inferred from conserved synteny, or seeded from a single query gene. We find SCoNe particularly helpful for exploring metagenomic dark matter, where genes are distant from database representatives and gene ontologies are often undescriptive. We release our SCoNe generation software at https://github.com/AaronAOliver/KiritimatiallesSCoNe to support its further use in visualization and hypothesis generation for meta-analyses and user-generated data.

## Acknowledgements

Computational analyses were performed using the San Diego Supercomputer Center’s Triton Shared Computing Cluster. This work was funded by United States Department of Agriculture Agriculture and Food Research Initiative Predoctoral Fellowship 2026-67011-46317 to A.O. and National Science Foundation grant OCE-2414798 and National Institutes of Health NIEHS grant P01ES035541-01 to E.E.A.

## Data Availability

Code used to generate SCoNes is available at https://github.com/AaronAOliver/KiritimatiallesSCoNe. SCoNes generated for Figures 3, 4, 5, and 6 are available as Cytoscape-compatible files on Zenodo 10.5281/zenodo.22103458. Metadata on genomes included in this study are included as **Supplemental Table 1**.

---

**Supplemental Figure 1.** P**o**lysaccharide **degradation potential varies by environment and taxonomic lineage.** (A) A principal component analysis of glycoside hydrolase, polysaccharide lyase, carbohydrate esterase, and sulfatase abundance in species representative *Kiritimatiellia* normalized by genes per genome. Points are individual genomes colored by environmental source. (B) The same principal components analysis plot, with *Pontiella* and CAIQIC01 colored as outliers.

**Supplemental Figure 2.** S**e**quence **similarity network of CAZyme gene cluster proteins in *Kiritimatiellia*.** Individual points are gene sequences from dbCAN-annotated CAZyme gene clusters, colored by putative function.

**Supplemental Figure 3.** B**i**osynthetic **potential of *Kiritimatiellia*.** Each bar graph shows the number of biosynthetic gene clusters in species representative *Kiritimatiellia* genomes as annotated by antiSMASH. Percentages display the ratio of each biosynthetic gene cluster classification to the total BGC set. The bottom axis uses a log scale.

**Supplemental Figure 4.** C**o**nserved **gene homology and synteny among *Pontiellaceae* and the wider *Kiritimatiellia*.** Homology map showing shared synteny blocks between *Pontiella desulfatans* and (A) other *Pontiellaceae* and (B) other *Kiritimatiellia*. Each line row represents one genome, and each column represents one gene in *P. desulfatans*. (C) The proportion of genomes that share a homolog with *P. desulfatans* at each protein position. Gray areas show synteny blocks explored in other figures.

