## Supplementary figures and images for "Conserved synteny reveals a saccharolytic lifestyle across *Kiritimatiellia*"

### Supplemental Figure 1

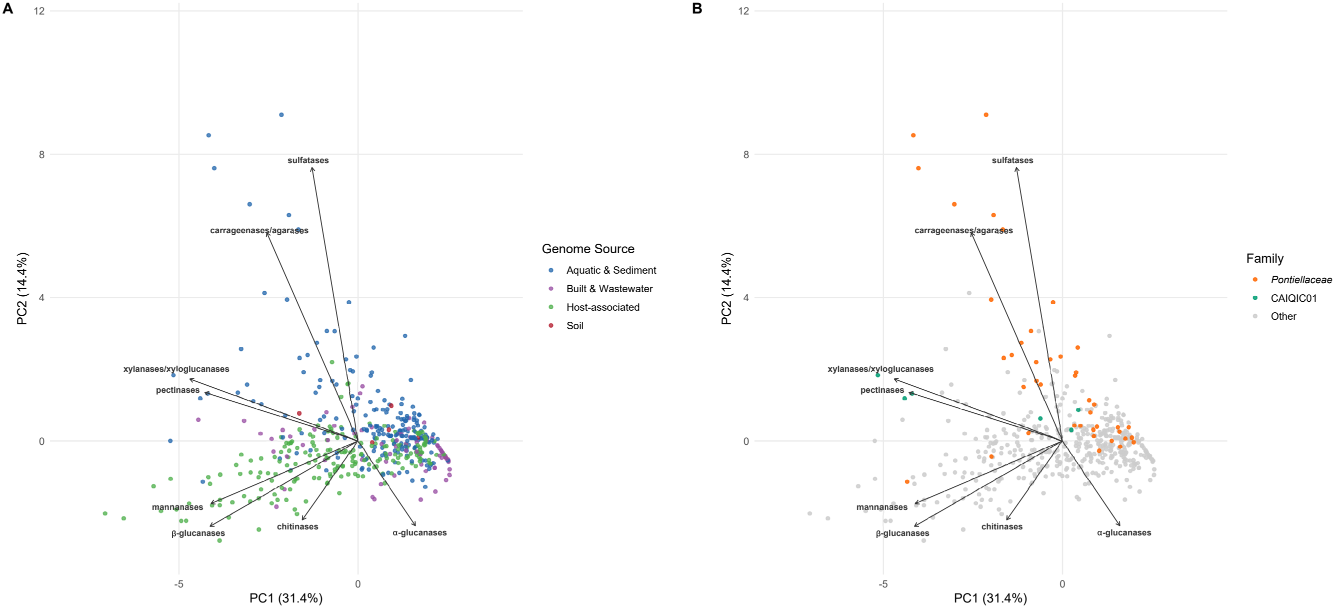

### Supplemental Figure 2

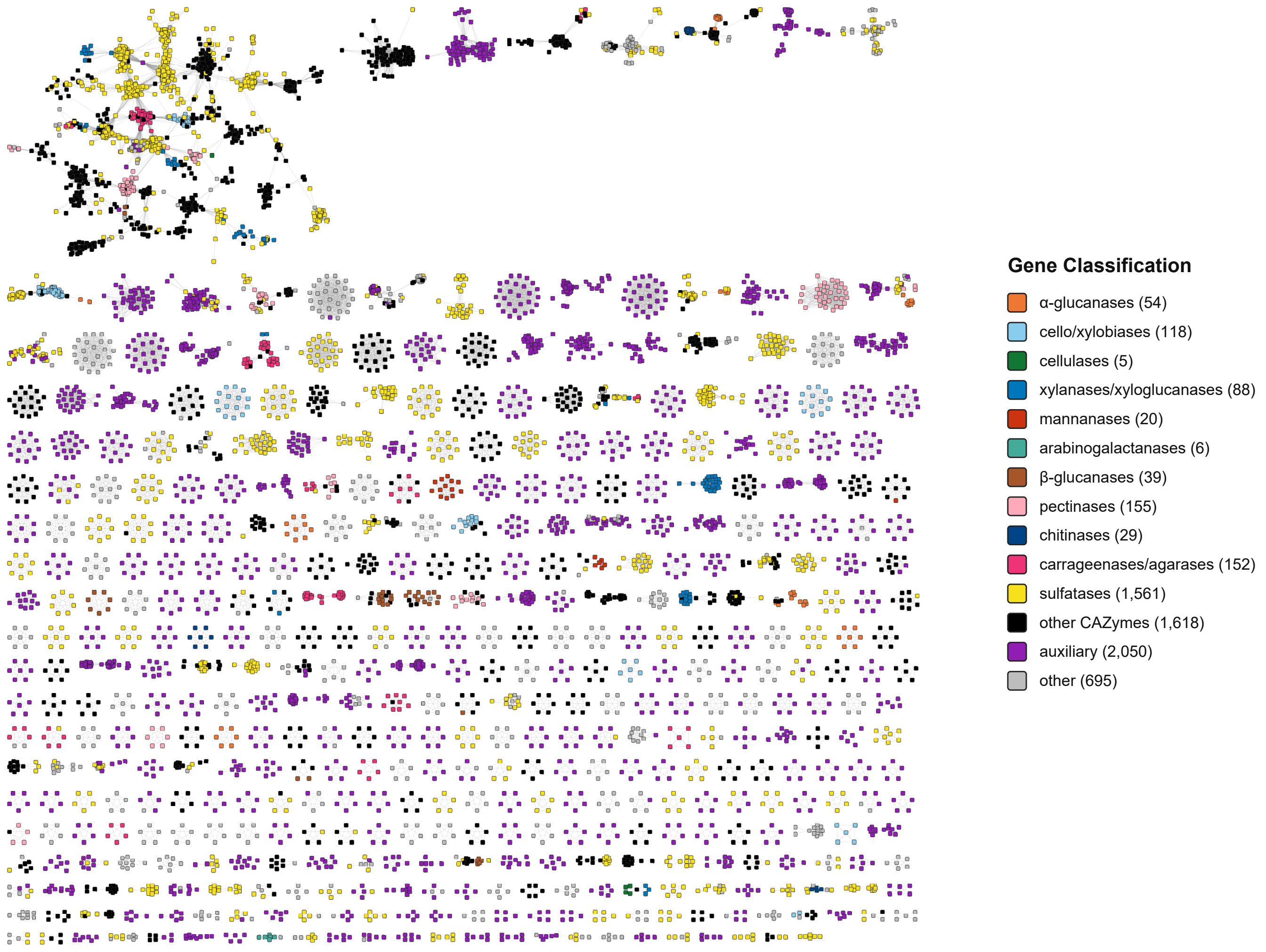

### Supplemental Figure 3

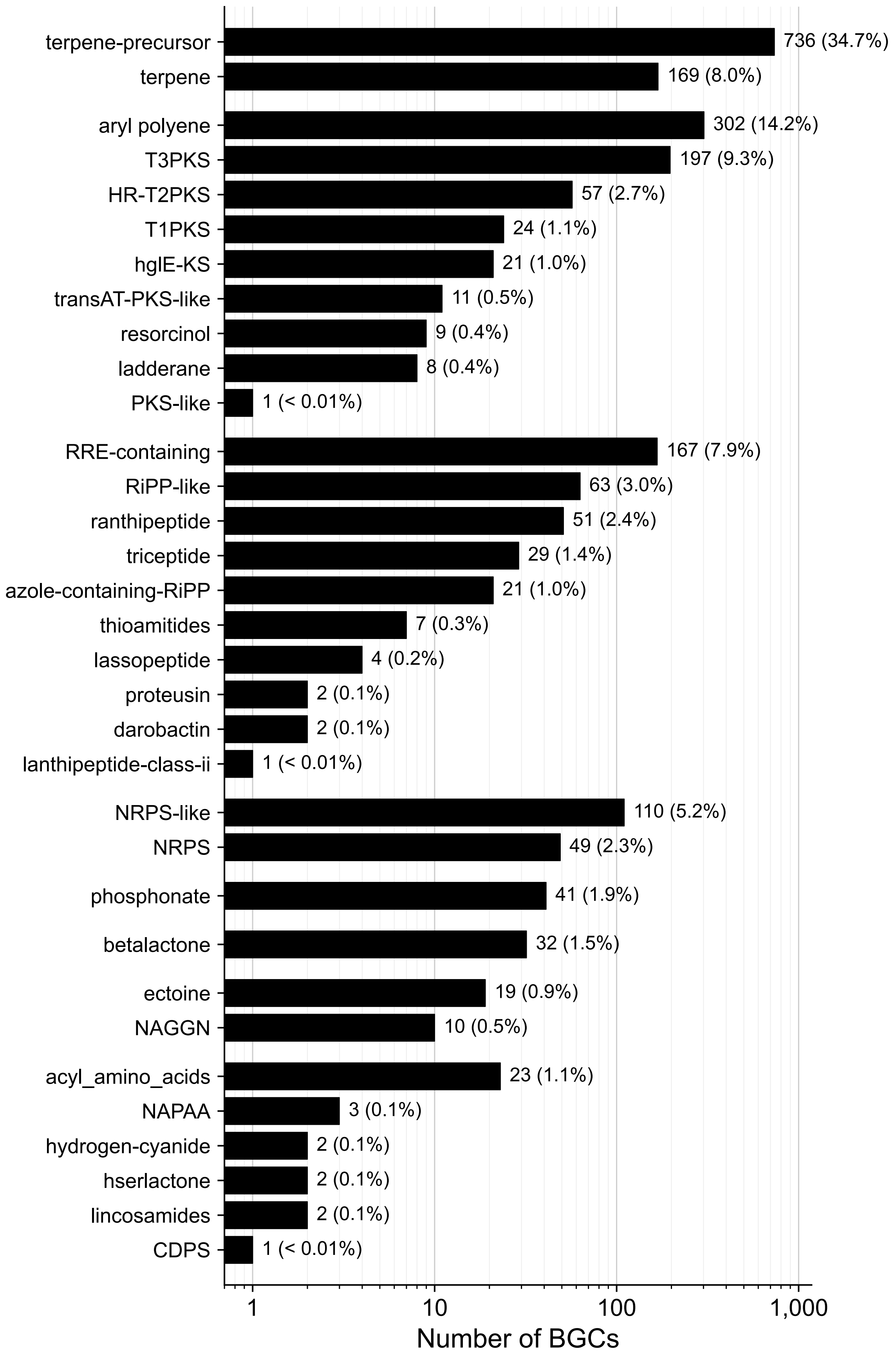

### Supplemental Figure 4

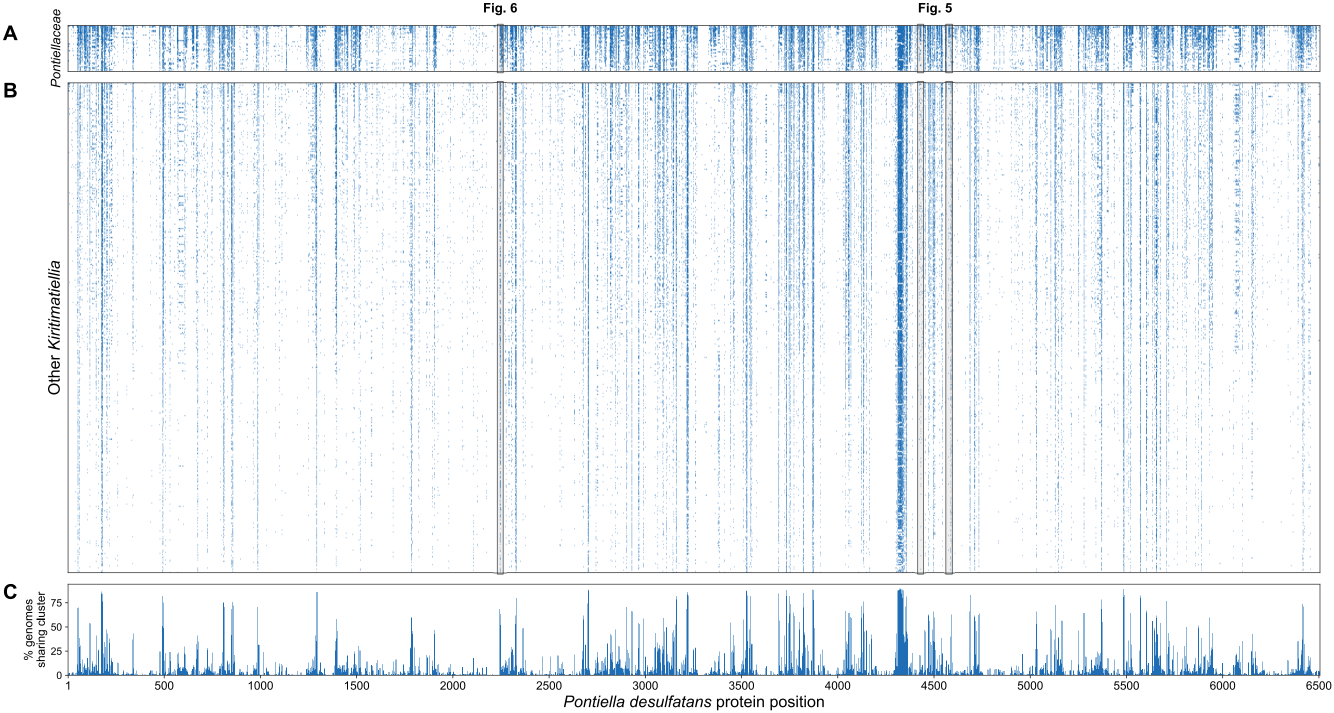
