## Supplemental Table 1 for "Conserved synteny reveals a saccharolytic lifestyle across *Kiritimatiellia*"

Supplemental Table 2.1: Metadata and taxonomy of species representative Kiritimatiellia genomes. The columns represent Genome Taxonomy Database accession, taxonomic strings for order and family, number of base pairs in the genome, number of coding sequences in the genome, and quality score calculated as completeness percentage – 5 x contamination percentage.

| GTDB Accession | GTDB Taxonomy | Genome Size | # CDS | Q-Score |
|---|---|---|---|---|
| GCA_003641465.1 | o__B70-G9;f__B70-G9 | 1147764 | 1109 | 55.94 |
| GCA_012961745.1 | o__B70-G9;f__B70-G9 | 1987671 | 1972 | 80.95 |
| GCA_029934055.1 | o__B70-G9;f__B70-G9 | 2200900 | 1860 | 84.49 |
| GCA_903824675.1 | o__CAIKKV01;f__CAIJXV01 | 4243228 | 3483 | 91.43 |
| GCA_945898015.1 | o__CAIKKV01;f__CAIKKV01 | 3292330 | 2590 | 64.99 |
| GCA_964201555.1 | o__CAIKKV01;f__CAIKKV01 | 1921903 | 1765 | 66.53 |
| GCA_938038445.1 | o__CAIKKV01;f__CAIKKV01 | 3069491 | 2441 | 91.94 |
| GCA_903895695.1 | o__CAIKKV01;f__CAITUY01 | 6341765 | 5170 | 83.16 |
| GCA_903908715.1 | o__CAIKKV01;f__CAITUY01 | 3052917 | 2692 | 83.75 |
| GCA_037128375.1 | o__CAIKKV01;f__CAITUY01 | 3638368 | 3632 | 73.94 |
| GCA_945870225.1 | o__CAIKKV01;f__CAITUY01 | 2702799 | 2243 | 71.78 |
| GCA_011391785.1 | o__CAIKKV01;f__CAITUY01 | 3066787 | 2717 | 82.42 |
| GCA_037128595.1 | o__CAIKKV01;f__CAITUY01 | 4680253 | 3915 | 91.12 |
| GCA_037131415.1 | o__CAIKKV01;f__CAITUY01 | 2808293 | 2816 | 67.88 |
| GCA_037132275.1 | o__CAIKKV01;f__CAITUY01 | 2669512 | 2398 | 86.19 |
| GCA_037134355.1 | o__CAIKKV01;f__CAITUY01 | 2965969 | 2783 | 81.18 |
| GCA_037141135.1 | o__CAIKKV01;f__CAITUY01 | 2931574 | 2755 | 93.74 |
| GCA_037143135.1 | o__CAIKKV01;f__CAITUY01 | 3382906 | 3300 | 81.51 |
| GCA_037143175.1 | o__CAIKKV01;f__CAITUY01 | 3303290 | 3084 | 87.42 |
| GCA_038330175.1 | o__CAIKKV01;f__CAITUY01 | 3774186 | 3427 | 77.42 |
| GCA_903934155.1 | o__CAIKKV01;f__CAITUY01 | 2515716 | 2429 | 71.8 |
| GCA_903958395.1 | o__CAIKKV01;f__CAITUY01 | 5202205 | 4337 | 85.22 |
| GCA_945890715.1 | o__CAIKKV01;f__CAITUY01 | 2106952 | 1953 | 83.06 |
| GCA_025360745.1 | o__CAIKKV01;f__CAIYGI01 | 2993897 | 2564 | 91.02 |
| GCA_037137675.1 | o__CAIKKV01;f__CAIYGI01 | 3253768 | 2988 | 74.46 |
| GCA_903925685.1 | o__CAIKKV01;f__CAIYGI01 | 4120073 | 3804 | 90.12 |
| GCA_938031185.1 | o__CAIKKV01;f__CALHGX01 | 2471062 | 2192 | 86.91 |
| GCA_026398385.1 | o__CAIQIC01;f__CAIQIC01 | 3661762 | 3355 | 76.58 |
| GCA_903871765.1 | o__CAIQIC01;f__CAIQIC01 | 3920532 | 3643 | 68.09 |
| GCA_004321905.1 | o__CAIQIC01;f__CAIQIC01 | 2529573 | 2267 | 93.18 |
| GCA_903889245.1 | o__CAIQIC01;f__CAIQIC01 | 5256829 | 4044 | 96.34 |
| GCA_903896925.1 | o__CAIQIC01;f__CAIQIC01 | 7224380 | 5341 | 91.55 |
| GCA_903912445.1 | o__CAIQIC01;f__CAIQIC01 | 6660355 | 5119 | 74.87 |
| GCA_001804865.1 | o__CAIQIC01;f__GWF2-57-35 | 5052682 | 3847 | 90.18 |
| GCA_937870865.1 | o__CAIQIC01;f__JAGNIJ01 | 2951872 | 2648 | 65.53 |

Supplemental Table 1: Metadata and taxonomy of species representative Kiritimatiellia genomes, Continued

| GTDB Accession | GTDB Taxonomy | Genome Size | # CDS | Q-Score |
|---|---|---|---|---|
| GCA_036821485.1 | o__CAIQIC01;f__JAGNIJ01 | 3650615 | 3055 | 84.72 |
| GCA_018001225.1 | o__CAIQIC01;f__JAGNIJ01 | 3963040 | 3140 | 92.85 |
| GCA_018823105.1 | o__CAIQIC01;f__JAGNIJ01 | 3564203 | 3175 | 85.06 |
| GCA_022710725.1 | o__CAIQIC01;f__JAGNIJ01 | 3886180 | 3011 | 93.63 |
| GCA_022704185.1 | o__CAIQIC01;f__JAGNIJ01 | 2407926 | 2213 | 66.43 |
| GCA_903906425.1 | o__CAIVKH01;f__CAIVKH01 | 6655884 | 5507 | 66.2 |
| GCA_035327305.1 | o__CAIVKH01;f__CAIVKH01 | 3272336 | 2716 | 92.58 |
| GCA_937872895.1 | o__CAIVKH01;f__CAIVKH01 | 3867962 | 3139 | 94.18 |
| GCA_937873455.1 | o__CAIVKH01;f__CAIVKH01 | 4121085 | 3338 | 90.04 |
| GCA_937894535.1 | o__CAIVKH01;f__CAIVKH01 | 3312000 | 3091 | 76.28 |
| GCA_035327665.1 | o__CAIVKH01;f__CAIVKH01 | 3627162 | 3070 | 92.61 |
| GCA_024233345.1 | o__CAIVKH01;f__CAIVKH01 | 3105184 | 2502 | 79.54 |
| GCA_023953695.1 | o__CAIVKH01;f__CAIVKH01 | 2620959 | 2305 | 93.09 |
| GCA_027493775.1 | o__CAIVKH01;f__CAIVKH01 | 3200265 | 3180 | 79.98 |
| GCA_035327585.1 | o__CAIVKH01;f__CAIVKH01 | 3799111 | 3138 | 94.37 |
| GCA_039807095.1 | o__CAIVKH01;f__CAIVKH01 | 2718640 | 2339 | 96.12 |
| GCA_937958885.1 | o__CAIVKH01;f__CAIVKH01 | 2580632 | 2294 | 87.72 |
| GCA_937996795.1 | o__CAIVKH01;f__CAIVKH01 | 3723485 | 3067 | 94.21 |
| GCA_025054615.1 | o__CAIVKH01;f__CAIVKH01 | 2482860 | 2155 | 93.87 |
| GCA_030432155.1 | o__CAIVKH01;f__CAIVKH01 | 3887255 | 3511 | 84.44 |
| GCA_913047795.1 | o__CAJWTS01;f__CAJWTS01 | 3497039 | 2987 | 85.68 |
| GCA_027623315.1 | o__CAJWTS01;f__CAJWTS01 | 3875074 | 3432 | 93.48 |
| GCA_964196025.1 | o__CAXMRT01;f__CAXMRT01 | 3403237 | 3086 | 73.89 |
| GCA_012514555.1 | o__JAAZAB01;f__JAAZAB01 | 3216425 | 2908 | 82.92 |
| GCA_016199105.1 | o__JAAZAB01;f__JAAZAB01 | 2913371 | 2806 | 56.39 |
| GCA_018898455.1 | o__JAAZAB01;f__JAAZAB01 | 3912958 | 3381 | 87.32 |
| GCA_020201415.1 | o__JAAZAB01;f__JAAZAB01 | 2688636 | 2695 | 61.33 |
| GCA_028715905.1 | o__JAAZAB01;f__JAAZAB01 | 2669920 | 2440 | 93.13 |
| GCA_041674125.1 | o__JAAZAB01;f__JAAZAB01 | 3876924 | 3318 | 75.89 |
| GCA_040767085.1 | o__JAAZAB01;f__JAAZAB01 | 4082876 | 3154 | 87.01 |
| GCA_040775225.1 | o__JAAZAB01;f__JAAZAB01 | 3851305 | 3259 | 74.16 |
| GCA_008636095.1 | o__JAAZAB01;f__JAAZAB01 | 3806325 | 3311 | 85.18 |
| GCA_016188985.1 | o__JAAZAB01;f__JAAZAB01 | 3085541 | 2641 | 80.98 |
| GCA_018896395.1 | o__JAAZAB01;f__JAAZAB01 | 2002574 | 1827 | 70.5 |
| GCA_026398415.1 | o__JAAZAB01;f__JAAZAB01 | 3294115 | 2762 | 91.03 |
| GCA_028718125.1 | o__JAAZAB01;f__JAAZAB01 | 4130942 | 3364 | 85.08 |
| GCA_041653875.1 | o__JAAZAB01;f__JAAZAB01 | 2690589 | 2330 | 90.84 |

Supplemental Table 1: Metadata and taxonomy of species representative Kiritimatiellia genomes, Continued

| GTDB Accession | GTDB Taxonomy | Genome Size | # CDS | Q-Score |
|---|---|---|---|---|
| GCA_016929905.1 | o__JAFGHX01;f__JAFGHX01 | 11135617 | 7700 | 64.65 |
| GCA_021734645.1 | o__JAIPIQ01;f__JAIPIQ01 | 2606670 | 2205 | 65.82 |
| GCA_029557905.1 | o__JAKQHR01;f__JAKQHR01 | 2548713 | 2374 | 81.7 |
| GCA_027858345.1 | o__JAQICX01;f__JAQICX01 | 4033302 | 3276 | 89.41 |
| GCA_030622955.1 | o__JAUXFM01;f__JAUXFM01 | 2652544 | 2483 | 70.12 |
| GCF_001017655.1 | o__Kiritimatiellales;f__Kiritimatiellaceae | 2949723 | 2429 | 97.02 |
| GCA_021734595.1 | o__Kiritimatiellales;f__Pontiellaceae | 3311578 | 2885 | 69.36 |
| GCF_034531395.1 | o__Kiritimatiellales;f__Pontiellaceae | 4436865 | 3611 | 94.29 |
| GCF_900890425.1 | o__Kiritimatiellales;f__Pontiellaceae | 8658305 | 6505 | 83.37 |
| GCA_029287275.1 | o__Kiritimatiellales;f__Pontiellaceae | 2866836 | 2876 | 73.09 |
| GCA_913065005.1 | o__Kiritimatiellales;f__Pontiellaceae | 1199496 | 1122 | 79.17 |
| GCA_002691095.1 | o__Kiritimatiellales;f__Pontiellaceae | 1303801 | 1199 | 63.17 |
| GCA_002712105.1 | o__Kiritimatiellales;f__Pontiellaceae | 1177491 | 1088 | 86.71 |
| GCA_003230915.1 | o__Kiritimatiellales;f__Pontiellaceae | 1903917 | 1882 | 86.68 |
| GCA_003246265.1 | o__Kiritimatiellales;f__Pontiellaceae | 2997214 | 2567 | 92.7 |
| GCA_003252225.1 | o__Kiritimatiellales;f__Pontiellaceae | 2999726 | 2851 | 81.18 |
| GCA_003532555.1 | o__Kiritimatiellales;f__Pontiellaceae | 1078159 | 1047 | 63.18 |
| GCA_004299845.1 | o__Kiritimatiellales;f__Pontiellaceae | 4107983 | 3666 | 84.49 |
| GCA_012729265.1 | o__Kiritimatiellales;f__Pontiellaceae | 3406929 | 3178 | 82.86 |
| GCA_013042165.1 | o__Kiritimatiellales;f__Pontiellaceae | 1851711 | 1942 | 68.23 |
| GCA_015486775.1 | o__Kiritimatiellales;f__Pontiellaceae | 2516066 | 2314 | 71.57 |
| GCA_016744085.1 | o__Kiritimatiellales;f__Pontiellaceae | 3004420 | 2910 | 91.35 |
| GCA_016937945.1 | o__Kiritimatiellales;f__Pontiellaceae | 4279068 | 3550 | 93.81 |
| GCA_021734345.1 | o__Kiritimatiellales;f__Pontiellaceae | 3247271 | 2773 | 73.33 |
| GCA_024279685.1 | o__Kiritimatiellales;f__Pontiellaceae | 2688584 | 2645 | 82.04 |
| GCA_029432775.1 | o__Kiritimatiellales;f__Pontiellaceae | 6294789 | 4872 | 88.81 |
| GCA_029432795.1 | o__Kiritimatiellales;f__Pontiellaceae | 4748661 | 4044 | 88.24 |
| GCA_029432815.1 | o__Kiritimatiellales;f__Pontiellaceae | 5377058 | 4483 | 90.32 |
| GCA_029935255.1 | o__Kiritimatiellales;f__Pontiellaceae | 3194711 | 3199 | 75.3 |
| GCA_029939385.1 | o__Kiritimatiellales;f__Pontiellaceae | 1742519 | 1822 | 57.94 |
| GCA_030595785.1 | o__Kiritimatiellales;f__Pontiellaceae | 1775158 | 1653 | 55.58 |
| GCA_030621805.1 | o__Kiritimatiellales;f__Pontiellaceae | 2414202 | 2446 | 65.92 |
| GCA_030625905.1 | o__Kiritimatiellales;f__Pontiellaceae | 2114843 | 2090 | 82.3 |
| GCA_030630445.1 | o__Kiritimatiellales;f__Pontiellaceae | 3394003 | 3159 | 77.74 |
| GCA_036479375.1 | o__Kiritimatiellales;f__Pontiellaceae | 2802826 | 2717 | 88.22 |
| GCA_036526075.1 | o__Kiritimatiellales;f__Pontiellaceae | 1545617 | 1505 | 83.15 |
| GCA_037132875.1 | o__Kiritimatiellales;f__Pontiellaceae | 3001690 | 2620 | 88.64 |

Supplemental Table 1: Metadata and taxonomy of species representative Kiritimatiellia genomes, Continued

| GTDB Accession | GTDB Taxonomy | Genome Size | # CDS | Q-Score |
|---|---|---|---|---|
| GCA_039714135.1 | o__Kiritimatiellales;f__Pontiellaceae | 2166254 | 2063 | 89.8 |
| GCA_039715135.1 | o__Kiritimatiellales;f__Pontiellaceae | 2652867 | 2513 | 67.31 |
| GCA_041582935.1 | o__Kiritimatiellales;f__Pontiellaceae | 3457235 | 2946 | 72.9 |
| GCF_041583075.1 | o__Kiritimatiellales;f__Pontiellaceae | 3606188 | 3242 | 73.54 |
| GCA_041592955.1 | o__Kiritimatiellales;f__Pontiellaceae | 2189613 | 2110 | 63.63 |
| GCA_041622415.1 | o__Kiritimatiellales;f__Pontiellaceae | 2880949 | 2759 | 61.15 |
| GCA_041740875.1 | o__Kiritimatiellales;f__Pontiellaceae | 3697861 | 3438 | 59.76 |
| GCA_937141895.1 | o__Kiritimatiellales;f__Pontiellaceae | 3059749 | 3228 | 52.79 |
| GCA_964014015.1 | o__Kiritimatiellales;f__Pontiellaceae | 1657538 | 1815 | 84.67 |
| GCA_964195195.1 | o__Kiritimatiellales;f__Pontiellaceae | 2193248 | 2305 | 65.39 |
| GCF_900890705.1 | o__Kiritimatiellales;f__Pontiellaceae | 7395795 | 5650 | 83.47 |
| GCF_009905215.1 | o__Kiritimatiellales;f__Tichowtungiaceae | 3882676 | 3157 | 92.83 |
| GCA_002423675.1 | o__Kiritimatiellales;f__Tichowtungiaceae | 1965982 | 2422 | 61.73 |
| GCA_013141415.1 | o__Kiritimatiellales;f__Tichowtungiaceae | 2218942 | 1938 | 94.45 |
| GCA_014381785.1 | o__Kiritimatiellales;f__Tichowtungiaceae | 2219109 | 2051 | 94.41 |
| GCA_016342445.1 | o__Kiritimatiellales;f__Tichowtungiaceae | 2085568 | 1955 | 82.22 |
| GCA_020853375.1 | o__Kiritimatiellales;f__Tichowtungiaceae | 1546500 | 1596 | 83.41 |
| GCA_025457785.1 | o__Kiritimatiellales;f__Tichowtungiaceae | 2464066 | 2335 | 77.68 |
| GCA_028692515.1 | o__Kiritimatiellales;f__Tichowtungiaceae | 1814136 | 1903 | 80.81 |
| GCA_029934215.1 | o__Kiritimatiellales;f__Tichowtungiaceae | 2737999 | 2457 | 86.25 |
| GCA_029939835.1 | o__Kiritimatiellales;f__Tichowtungiaceae | 1927440 | 1957 | 84.37 |
| GCA_030153935.1 | o__Kiritimatiellales;f__Tichowtungiaceae | 2035175 | 1963 | 91.33 |
| GCA_030619095.1 | o__Kiritimatiellales;f__Tichowtungiaceae | 1573560 | 1733 | 72.04 |
| GCA_035404105.1 | o__Kiritimatiellales;f__Tichowtungiaceae | 1819721 | 1745 | 89.43 |
| GCA_037141075.1 | o__Kiritimatiellales;f__Tichowtungiaceae | 1997814 | 1930 | 80.77 |
| GCA_037144035.1 | o__Kiritimatiellales;f__Tichowtungiaceae | 2399162 | 2441 | 75.99 |
| GCA_038213895.1 | o__Kiritimatiellales;f__Tichowtungiaceae | 3154505 | 2989 | 78.88 |
| GCA_041584215.1 | o__Kiritimatiellales;f__Tichowtungiaceae | 4501659 | 3448 | 77.86 |
| GCA_041595735.1 | o__Kiritimatiellales;f__Tichowtungiaceae | 3244441 | 2832 | 78.56 |
| GCA_041656295.1 | o__Kiritimatiellales;f__Tichowtungiaceae | 2859210 | 2578 | 98.13 |
| GCA_041661845.1 | o__Kiritimatiellales;f__Tichowtungiaceae | 3354091 | 2973 | 79.26 |
| GCA_041674365.1 | o__Kiritimatiellales;f__Tichowtungiaceae | 1862000 | 2015 | 76.19 |
| GCA_903826865.1 | o__Kiritimatiellales;f__Tichowtungiaceae | 2644415 | 2374 | 97.42 |
| GCA_903831085.1 | o__Kiritimatiellales;f__Tichowtungiaceae | 2138574 | 1936 | 77.06 |
| GCA_017527155.1 | o__LD1-PB3_B;f__Lenti-01 | 3227041 | 2631 | 92.6 |
| GCA_017540905.1 | o__LD1-PB3_B;f__Lenti-01 | 3507924 | 2825 | 64.86 |
| GCA_902766695.1 | o__LD1-PB3_B;f__Lenti-01 | 3036038 | 2564 | 91.96 |

Supplemental Table 1: Metadata and taxonomy of species representative Kiritimatiellia genomes, Continued

| GTDB Accession | GTDB Taxonomy | Genome Size | # CDS | Q-Score |
|---|---|---|---|---|
| GCA_012511885.1 | o__LD1-PB3_B;f__Lenti-01 | 2494825 | 1990 | 84.35 |
| GCA_018432565.1 | o__LD1-PB3_B;f__Lenti-01 | 2508724 | 2502 | 74.65 |
| GCA_028681135.1 | o__LD1-PB3_B;f__Lenti-01 | 3048864 | 2524 | 93.16 |
| GCA_034005635.1 | o__LD1-PB3_B;f__Lenti-01 | 2458258 | 2133 | 86.39 |
| GCA_040768005.1 | o__LD1-PB3_B;f__Lenti-01 | 2576830 | 2217 | 82.39 |
| GCA_031262805.1 | o__LD1-PB3_B;f__Lenti-01 | 1981691 | 1764 | 82.66 |
| GCA_002071335.1 | o__LD1-PB3_B;f__Lenti-01 | 2752870 | 2643 | 75.43 |
| GCA_002304915.1 | o__LD1-PB3_B;f__Lenti-01 | 2108534 | 1859 | 85.72 |
| GCA_002347655.1 | o__LD1-PB3_B;f__Lenti-01 | 3105763 | 2580 | 96.2 |
| GCA_003451935.1 | o__LD1-PB3_B;f__Lenti-01 | 1850317 | 1828 | 52.83 |
| GCA_009928955.1 | o__LD1-PB3_B;f__Lenti-01 | 3423791 | 2961 | 50.45 |
| GCA_009930415.1 | o__LD1-PB3_B;f__Lenti-01 | 3104077 | 2842 | 88.61 |
| GCA_012514035.1 | o__LD1-PB3_B;f__Lenti-01 | 2784004 | 2357 | 88.92 |
| GCA_022709625.1 | o__LD1-PB3_B;f__Lenti-01 | 2259818 | 2273 | 64.78 |
| GCA_035450765.1 | o__LD1-PB3_B;f__Lenti-01 | 2875696 | 2750 | 84.18 |
| GCA_937860055.1 | o__LD1-PB3_B;f__Lenti-01 | 3049884 | 2502 | 92.31 |
| GCA_937872095.1 | o__LD1-PB3_B;f__Lenti-01 | 3331579 | 2700 | 87.82 |
| GCA_938003985.1 | o__LD1-PB3_B;f__Lenti-01 | 3118850 | 2592 | 95.22 |
| GCA_017445365.1 | o__LD1-PB3_B;f__Lenti-01 | 2325113 | 2189 | 62.26 |
| GCA_017458625.1 | o__LD1-PB3_B;f__Lenti-01 | 2691885 | 2174 | 78.59 |
| GCA_026417735.1 | o__PWTM01;f__PWTM01 | 3094483 | 2484 | 95.3 |
| GCA_900661335.1 | o__PWTM01;f__PWTM01 | 2632675 | 2178 | 67.15 |
| GCA_937139365.1 | o__PWTM01;f__PWTM01 | 2825592 | 2705 | 93.65 |
| GCA_938010045.1 | o__PWTM01;f__PWTM01 | 3215806 | 2647 | 97.27 |
| GCA_937935255.1 | o__PWTM01;f__PWTM01 | 2734788 | 2316 | 85.6 |
| GCA_938005705.1 | o__PWTM01;f__PWTM01 | 2756372 | 2499 | 90.51 |
| GCA_017996315.1 | o__PWTM01;f__PWTM01 | 2221561 | 2021 | 69.69 |
| GCA_003563855.1 | o__PWTM01;f__PWTM01 | 3274399 | 2838 | 88.41 |
| GCA_016873715.1 | o__PWTM01;f__PWTM01 | 5126434 | 3944 | 89.04 |
| GCA_003565195.1 | o__PWZG01;f__PXCX01 | 2385145 | 2175 | 80.22 |
| GCA_003567575.1 | o__PWZG01;f__PXCX01 | 4034550 | 3450 | 72.13 |
| GCA_037129105.1 | o__RFP12;f__CAIWXQ01 | 3104743 | 2608 | 71.24 |
| GCA_037144015.1 | o__RFP12;f__CAIWXQ01 | 3224386 | 2598 | 89.01 |
| GCA_903861045.1 | o__RFP12;f__CAIWXQ01 | 4726601 | 3781 | 85.65 |
| GCA_903916675.1 | o__RFP12;f__CAIWXQ01 | 2477839 | 2020 | 86.52 |
| GCA_012514915.1 | o__RFP12;f__JAAYLN01 | 2778191 | 2604 | 65.58 |
| GCA_012521875.1 | o__RFP12;f__JAAYLN01 | 3955686 | 3106 | 85.22 |

Supplemental Table 1: Metadata and taxonomy of species representative Kiritimatiellia genomes, Continued

| GTDB Accession | GTDB Taxonomy | Genome Size | # CDS | Q-Score |
|---|---|---|---|---|
| GCA_028718445.1 | o__RFP12;f__JAAYLN01 | 5721928 | 4809 | 58.48 |
| GCA_017458965.1 | o__RFP12;f__JAAYLN01 | 1750765 | 1422 | 57.11 |
| GCA_017509565.1 | o__RFP12;f__JAAYLN01 | 3197410 | 2697 | 85.58 |
| GCA_017962605.1 | o__RFP12;f__JAAYLN01 | 4181991 | 3248 | 76.37 |
| GCA_012520735.1 | o__RFP12;f__JAAYNR01 | 2765688 | 2363 | 89.25 |
| GCA_016873675.1 | o__RFP12;f__JAAYNR01 | 3169171 | 2633 | 92.28 |
| GCA_945870025.1 | o__RFP12;f__JAAYNR01 | 2755496 | 2105 | 81.26 |
| GCA_031550185.1 | o__RFP12;f__JAVBOU01 | 2947566 | 2299 | 87.34 |
| GCA_902773475.1 | o__RFP12;f__UBA1067 | 1782564 | 1639 | 71.91 |
| GCA_902799715.1 | o__RFP12;f__UBA1067 | 2551216 | 2009 | 89.35 |
| GCA_037133395.1 | o__RFP12;f__UBA1067 | 5037336 | 3708 | 80.55 |
| GCA_037141675.1 | o__RFP12;f__UBA1067 | 4258583 | 3697 | 77.77 |
| GCA_903935195.1 | o__RFP12;f__UBA1067 | 3626931 | 3211 | 83.31 |
| GCA_945870145.1 | o__RFP12;f__UBA1067 | 2433651 | 2128 | 71.17 |
| GCA_946623785.1 | o__RFP12;f__UBA1067 | 2790471 | 2276 | 55.29 |
| GCA_035353175.1 | o__RFP12;f__UBA1067 | 5319760 | 3745 | 89.4 |
| GCA_002071025.1 | o__RFP12;f__UBA1067 | 4443224 | 3934 | 71.72 |
| GCA_012838435.1 | o__RFP12;f__UBA1067 | 3998345 | 3196 | 82.1 |
| GCA_017999015.1 | o__RFP12;f__UBA1067 | 3558442 | 3028 | 76.03 |
| GCA_022704125.1 | o__RFP12;f__UBA1067 | 2469823 | 2014 | 55.6 |
| GCA_028684935.1 | o__RFP12;f__UBA1067 | 3807257 | 2982 | 81.34 |
| GCA_035449085.1 | o__RFP12;f__UBA1067 | 3271683 | 2458 | 74.84 |
| GCA_035450385.1 | o__RFP12;f__UBA1067 | 3500488 | 2729 | 60.13 |
| GCA_023269675.1 | o__RFP12;f__UBA1067 | 6325037 | 4298 | 92.16 |
| GCA_025360845.1 | o__RFP12;f__UBA1067 | 5169291 | 4108 | 89.1 |
| GCA_038219365.1 | o__RFP12;f__UBA1067 | 3078675 | 2659 | 72.94 |
| GCA_022704145.1 | o__RFP12;f__UBA1067 | 4570006 | 3552 | 68.17 |
| GCA_028700415.1 | o__RFP12;f__UBA1067 | 4912576 | 3937 | 76.36 |
| GCA_033812795.1 | o__RFP12;f__UBA1067 | 3537586 | 2723 | 90.68 |
| GCA_034003585.1 | o__RFP12;f__UBA1067 | 3732651 | 2830 | 94.72 |
| GCA_028701555.1 | o__RFP12;f__UBA1067 | 2690490 | 2503 | 66.87 |
| GCA_028707675.1 | o__RFP12;f__UBA1067 | 2284101 | 2247 | 73.65 |
| GCA_031553315.1 | o__RFP12;f__UBA1067 | 2469771 | 1947 | 94 |
| GCA_031552845.1 | o__RFP12;f__UBA1067 | 2982716 | 2378 | 92.32 |
| GCA_034178815.1 | o__RFP12;f__UBA1067 | 2020783 | 1756 | 90.85 |
| GCA_018399555.1 | o__RFP12;f__UBA1067 | 2860232 | 2262 | 81.13 |
| GCA_027858965.1 | o__RFP12;f__UBA1067 | 5931720 | 4428 | 88.09 |

Supplemental Table 1: Metadata and taxonomy of species representative Kiritimatiellia genomes, Continued

| GTDB Accession | GTDB Taxonomy | Genome Size | # CDS | Q-Score |
|---|---|---|---|---|
| GCA_028686345.1 | o__RFP12;f__UBA1067 | 5298950 | 4044 | 83.99 |
| GCA_041673345.1 | o__RFP12;f__UBA1067 | 4630186 | 4056 | 64.3 |
| GCA_017479035.1 | o__RFP12;f__UBA1067 | 1649954 | 1467 | 66.28 |
| GCA_017525225.1 | o__RFP12;f__UBA1067 | 4201823 | 3292 | 87.35 |
| GCA_024405395.1 | o__RFP12;f__UBA1067 | 4044349 | 2936 | 77.35 |
| GCA_017450695.1 | o__RFP12;f__UBA1067 | 5025092 | 3593 | 81.99 |
| GCA_017544725.1 | o__RFP12;f__UBA1067 | 3803722 | 2847 | 61.67 |
| GCA_017613085.1 | o__RFP12;f__UBA1067 | 2426471 | 1992 | 65.29 |
| GCA_017962315.1 | o__RFP12;f__UBA1067 | 3261983 | 2544 | 81.9 |
| GCA_017646325.1 | o__RFP12;f__UBA1067 | 1397980 | 1246 | 60.62 |
| GCA_002373695.1 | o__RFP12;f__UBA1067 | 3529299 | 2663 | 78.06 |
| GCA_016302985.1 | o__RFP12;f__UBA1067 | 3267215 | 2492 | 63.88 |
| GCA_017411125.1 | o__RFP12;f__UBA1067 | 2746143 | 2496 | 75.17 |
| GCA_017427485.1 | o__RFP12;f__UBA1067 | 6301721 | 4695 | 55.01 |
| GCA_017430155.1 | o__RFP12;f__UBA1067 | 3875608 | 3326 | 61.1 |
| GCA_017459225.1 | o__RFP12;f__UBA1067 | 3676726 | 3033 | 82.13 |
| GCA_017563185.1 | o__RFP12;f__UBA1067 | 3207387 | 2520 | 92.82 |
| GCA_030537465.1 | o__RFP12;f__UBA1067 | 5188241 | 3858 | 74.55 |
| GCA_900320225.1 | o__RFP12;f__UBA1067 | 5214994 | 4177 | 85.9 |
| GCA_900547945.1 | o__RFP12;f__UBA1067 | 3452514 | 2664 | 90.46 |
| GCA_900771305.1 | o__RFP12;f__UBA1067 | 3810265 | 3253 | 80.98 |
| GCA_902761035.1 | o__RFP12;f__UBA1067 | 6137606 | 4816 | 74.7 |
| GCA_902763655.1 | o__RFP12;f__UBA1067 | 4502110 | 3671 | 83.09 |
| GCA_902769585.1 | o__RFP12;f__UBA1067 | 4903547 | 3907 | 53.69 |
| GCA_902779865.1 | o__RFP12;f__UBA1067 | 4298331 | 3474 | 76.05 |
| GCA_902802945.1 | o__RFP12;f__UBA1067 | 5813896 | 4506 | 74.2 |
| GCA_902803135.1 | o__RFP12;f__UBA1067 | 3114024 | 2438 | 78.17 |
| GCA_934643865.1 | o__RFP12;f__UBA1067 | 3308636 | 2440 | 92.98 |
| GCA_934667295.1 | o__RFP12;f__UBA1067 | 1764331 | 1577 | 76.92 |
| GCA_946401765.1 | o__RFP12;f__UBA1067 | 3179493 | 2552 | 67.23 |
| GCA_946413265.1 | o__RFP12;f__UBA1067 | 4701272 | 3610 | 87.37 |
| GCA_946543025.1 | o__RFP12;f__UBA1067 | 3565813 | 2749 | 72.48 |
| GCA_963614205.1 | o__RFP12;f__UBA1067 | 1755976 | 1546 | 74.91 |
| GCA_963618935.1 | o__RFP12;f__UBA1067 | 2644887 | 2254 | 80.26 |
| GCA_963629855.1 | o__RFP12;f__UBA1067 | 1285335 | 1327 | 53.61 |
| GCA_018713405.1 | o__RFP12;f__UBA1067 | 1778802 | 1624 | 86.45 |
| GCA_004555175.1 | o__RFP12;f__UBA1067 | 2078870 | 1614 | 92.72 |

Supplemental Table 1: Metadata and taxonomy of species representative Kiritimatiellia genomes, Continued

| GTDB Accession | GTDB Taxonomy | Genome Size | # CDS | Q-Score |
|---|---|---|---|---|
| GCA_017457555.1 | o__RFP12;f__UBA1067 | 1913838 | 1775 | 81.95 |
| GCA_017531445.1 | o__RFP12;f__UBA1067 | 1404358 | 1122 | 49.86 |
| GCA_017548245.1 | o__RFP12;f__UBA1067 | 1390891 | 1341 | 74.82 |
| GCA_017558005.1 | o__RFP12;f__UBA1067 | 2051466 | 1587 | 88.97 |
| GCA_017646485.1 | o__RFP12;f__UBA1067 | 1482576 | 1420 | 60.08 |
| GCA_030535915.1 | o__RFP12;f__UBA1067 | 1673446 | 1446 | 86.96 |
| GCA_036651955.1 | o__RFP12;f__UBA1067 | 1699609 | 1450 | 90.43 |
| GCA_900771325.1 | o__RFP12;f__UBA1067 | 2426387 | 1890 | 80.76 |
| GCA_937892925.1 | o__RFP12;f__UBA1067 | 2005506 | 1681 | 94.4 |
| GCA_944389545.1 | o__RFP12;f__UBA1067 | 2534065 | 1834 | 95.62 |
| GCA_944389565.1 | o__RFP12;f__UBA1067 | 2256776 | 1817 | 94.64 |
| GCA_944389585.1 | o__RFP12;f__UBA1067 | 1949089 | 1595 | 93.03 |
| GCA_944389945.1 | o__RFP12;f__UBA1067 | 2325100 | 1783 | 90.45 |
| GCA_944391055.1 | o__RFP12;f__UBA1067 | 2153977 | 1748 | 93.4 |
| GCA_944391065.1 | o__RFP12;f__UBA1067 | 1956537 | 1609 | 87.93 |
| GCA_944391085.1 | o__RFP12;f__UBA1067 | 1727642 | 1437 | 88.56 |
| GCA_946056345.1 | o__RFP12;f__UBA1067 | 1445761 | 1305 | 74.79 |
| GCA_949282555.1 | o__RFP12;f__UBA1067 | 2426525 | 1963 | 93.95 |
| GCA_949288955.1 | o__RFP12;f__UBA1067 | 2248286 | 1790 | 91.79 |
| GCA_949289005.1 | o__RFP12;f__UBA1067 | 2357542 | 1901 | 93.91 |
| GCA_963608725.1 | o__RFP12;f__UBA1067 | 1671341 | 1387 | 93.39 |
| GCA_002298665.1 | o__RFP12;f__UBA1067 | 1553145 | 1373 | 90.14 |
| GCA_002315875.1 | o__RFP12;f__UBA1067 | 2317670 | 1923 | 54.07 |
| GCA_002321305.1 | o__RFP12;f__UBA1067 | 2723985 | 2316 | 90.03 |
| GCA_902775595.1 | o__RFP12;f__UBA1067 | 3153127 | 2556 | 72.4 |
| GCA_902764415.1 | o__RFP12;f__UBA1067 | 2599316 | 2073 | 88.14 |
| GCA_946613265.1 | o__RFP12;f__UBA1067 | 2229349 | 1741 | 80.02 |
| GCA_002369235.1 | o__RFP12;f__UBA1067 | 1639202 | 1494 | 85.86 |
| GCA_002371135.1 | o__RFP12;f__UBA1067 | 2589698 | 2022 | 83.25 |
| GCA_002371755.1 | o__RFP12;f__UBA1067 | 2427939 | 2089 | 73.92 |
| GCA_002392765.1 | o__RFP12;f__UBA1067 | 3047170 | 2387 | 83.45 |
| GCA_946544115.1 | o__RFP12;f__UBA1067 | 2282582 | 1866 | 66.39 |
| GCA_002440145.1 | o__RFP12;f__UBA1067 | 1732897 | 1519 | 85.02 |
| GCA_946533675.1 | o__RFP12;f__UBA1067 | 2608011 | 2000 | 89.3 |
| GCA_004551905.1 | o__RFP12;f__UBA1067 | 2004189 | 1692 | 76.55 |
| GCA_015061665.1 | o__RFP12;f__UBA1067 | 3440937 | 2756 | 89.2 |
| GCA_016280825.1 | o__RFP12;f__UBA1067 | 2318254 | 2048 | 59.31 |

Supplemental Table 1: Metadata and taxonomy of species representative Kiritimatiellia genomes, Continued

| GTDB Accession | GTDB Taxonomy | Genome Size | # CDS | Q-Score |
|---|---|---|---|---|
| GCA_016284515.1 | o__RFP12;f__UBA1067 | 3860512 | 2930 | 76.3 |
| GCA_016288465.1 | o__RFP12;f__UBA1067 | 3524107 | 2830 | 60.42 |
| GCA_016296065.1 | o__RFP12;f__UBA1067 | 3703418 | 2686 | 73.15 |
| GCA_016303225.1 | o__RFP12;f__UBA1067 | 2897247 | 2485 | 50.92 |
| GCA_017385345.1 | o__RFP12;f__UBA1067 | 1571623 | 1579 | 57.51 |
| GCA_017395505.1 | o__RFP12;f__UBA1067 | 1817480 | 1716 | 90.05 |
| GCA_017403825.1 | o__RFP12;f__UBA1067 | 3932509 | 3026 | 71.39 |
| GCA_017406465.1 | o__RFP12;f__UBA1067 | 3506975 | 2721 | 71.84 |
| GCA_017412325.1 | o__RFP12;f__UBA1067 | 878340 | 952 | 53.64 |
| GCA_017418445.1 | o__RFP12;f__UBA1067 | 1386563 | 1229 | 63.66 |
| GCA_017432385.1 | o__RFP12;f__UBA1067 | 2736638 | 2370 | 65.38 |
| GCA_017447805.1 | o__RFP12;f__UBA1067 | 2248606 | 2098 | 84.75 |
| GCA_017463935.1 | o__RFP12;f__UBA1067 | 2794465 | 2628 | 79.55 |
| GCA_017507065.1 | o__RFP12;f__UBA1067 | 1249332 | 1188 | 68.48 |
| GCA_017508315.1 | o__RFP12;f__UBA1067 | 2881069 | 2332 | 76 |
| GCA_017514235.1 | o__RFP12;f__UBA1067 | 3959046 | 3139 | 83.56 |
| GCA_017520845.1 | o__RFP12;f__UBA1067 | 2740765 | 2160 | 89.7 |
| GCA_017525475.1 | o__RFP12;f__UBA1067 | 1315052 | 1334 | 81.39 |
| GCA_017542055.1 | o__RFP12;f__UBA1067 | 1346471 | 1295 | 88.53 |
| GCA_017628025.1 | o__RFP12;f__UBA1067 | 1448485 | 1245 | 61.27 |
| GCA_017628165.1 | o__RFP12;f__UBA1067 | 1238172 | 1178 | 60.46 |
| GCA_017629735.1 | o__RFP12;f__UBA1067 | 1480835 | 1375 | 62.7 |
| GCA_017631385.1 | o__RFP12;f__UBA1067 | 2019777 | 1812 | 70.29 |
| GCA_017648225.1 | o__RFP12;f__UBA1067 | 1876072 | 1810 | 71.4 |
| GCA_017650345.1 | o__RFP12;f__UBA1067 | 1442321 | 1248 | 64.69 |
| GCA_017652705.1 | o__RFP12;f__UBA1067 | 1660893 | 1418 | 62.09 |
| GCA_017959645.1 | o__RFP12;f__UBA1067 | 1974595 | 1835 | 60.78 |
| GCA_017965685.1 | o__RFP12;f__UBA1067 | 1411607 | 1437 | 61.73 |
| GCA_018384485.1 | o__RFP12;f__UBA1067 | 1819028 | 1561 | 79.64 |
| GCA_024405265.1 | o__RFP12;f__UBA1067 | 1707718 | 1499 | 77.1 |
| GCA_024694365.1 | o__RFP12;f__UBA1067 | 2461998 | 2099 | 58.97 |
| GCA_900320855.1 | o__RFP12;f__UBA1067 | 2409262 | 2004 | 80.91 |
| GCA_900321255.1 | o__RFP12;f__UBA1067 | 2776193 | 2364 | 81.48 |
| GCA_900321575.1 | o__RFP12;f__UBA1067 | 1439611 | 1345 | 75 |
| GCA_900767325.1 | o__RFP12;f__UBA1067 | 1101788 | 1016 | 58.16 |
| GCA_902761275.1 | o__RFP12;f__UBA1067 | 3425386 | 2886 | 74.54 |
| GCA_902770045.1 | o__RFP12;f__UBA1067 | 2270887 | 1868 | 83.71 |

Supplemental Table 1: Metadata and taxonomy of species representative Kiritimatiellia genomes, Continued

| GTDB Accession | GTDB Taxonomy | Genome Size | # CDS | Q-Score |
|---|---|---|---|---|
| GCA_902772905.1 | o__RFP12;f__UBA1067 | 2360034 | 2144 | 68.21 |
| GCA_902773015.1 | o__RFP12;f__UBA1067 | 2240586 | 1959 | 64.83 |
| GCA_902776615.1 | o__RFP12;f__UBA1067 | 2354932 | 2082 | 80.6 |
| GCA_902780865.1 | o__RFP12;f__UBA1067 | 3356427 | 2709 | 64.12 |
| GCA_902788415.1 | o__RFP12;f__UBA1067 | 1897223 | 1711 | 76.02 |
| GCA_902799385.1 | o__RFP12;f__UBA1067 | 2708262 | 2373 | 71.05 |
| GCA_902799955.1 | o__RFP12;f__UBA1067 | 3808006 | 2938 | 66.58 |
| GCA_902800055.1 | o__RFP12;f__UBA1067 | 2286154 | 1869 | 85.95 |
| GCA_902801025.1 | o__RFP12;f__UBA1067 | 3501112 | 2809 | 65.83 |
| GCA_904502135.1 | o__RFP12;f__UBA1067 | 1921602 | 1709 | 86.86 |
| GCA_934621385.1 | o__RFP12;f__UBA1067 | 2317810 | 1962 | 84.24 |
| GCA_934635045.1 | o__RFP12;f__UBA1067 | 3688694 | 2708 | 89.1 |
| GCA_934635225.1 | o__RFP12;f__UBA1067 | 2943281 | 2315 | 89.47 |
| GCA_934635385.1 | o__RFP12;f__UBA1067 | 3933021 | 3161 | 81.67 |
| GCA_934654805.1 | o__RFP12;f__UBA1067 | 3174523 | 2508 | 89.35 |
| GCA_934675625.1 | o__RFP12;f__UBA1067 | 3593279 | 2777 | 88.74 |
| GCA_937901395.1 | o__RFP12;f__UBA1067 | 1549108 | 1465 | 65.93 |
| GCA_945487985.1 | o__RFP12;f__UBA1067 | 1472395 | 1324 | 60.53 |
| GCA_945613785.1 | o__RFP12;f__UBA1067 | 1764002 | 1660 | 67.38 |
| GCA_945834475.1 | o__RFP12;f__UBA1067 | 2567673 | 2226 | 82.94 |
| GCA_945947925.1 | o__RFP12;f__UBA1067 | 2042908 | 1824 | 73.39 |
| GCA_946055075.1 | o__RFP12;f__UBA1067 | 1996835 | 1757 | 80.65 |
| GCA_946610035.1 | o__RFP12;f__UBA1067 | 3033296 | 2309 | 77.59 |
| GCA_946615875.1 | o__RFP12;f__UBA1067 | 1743495 | 1465 | 53.93 |
| GCA_946628745.1 | o__RFP12;f__UBA1067 | 1558805 | 1417 | 54.23 |
| GCA_947305585.1 | o__RFP12;f__UBA1067 | 1159760 | 1168 | 64.38 |
| GCA_947305775.1 | o__RFP12;f__UBA1067 | 2702525 | 2350 | 83.69 |
| GCA_963588415.1 | o__RFP12;f__UBA1067 | 2303474 | 2022 | 86.14 |
| GCA_963600185.1 | o__RFP12;f__UBA1067 | 1987143 | 1679 | 83.05 |
| GCA_963602475.1 | o__RFP12;f__UBA1067 | 1946191 | 1633 | 88.18 |
| GCA_963606275.1 | o__RFP12;f__UBA1067 | 1416086 | 1368 | 56.93 |
| GCA_963612485.1 | o__RFP12;f__UBA1067 | 1898984 | 1751 | 79.96 |
| GCA_963616865.1 | o__RFP12;f__UBA1067 | 1152615 | 1155 | 65.81 |
| GCA_963626175.1 | o__RFP12;f__UBA1067 | 2049293 | 1890 | 65.77 |
| GCA_963629255.1 | o__RFP12;f__UBA1067 | 2205502 | 1937 | 87.54 |
| GCA_963630185.1 | o__RFP12;f__UBA1067 | 1597550 | 1509 | 64.31 |
| GCA_963636335.1 | o__RFP12;f__UBA1067 | 1449512 | 1382 | 81.18 |

Supplemental Table 1: Metadata and taxonomy of species representative Kiritimatiellia genomes, Continued

| GTDB Accession | GTDB Taxonomy | Genome Size | # CDS | Q-Score |
|---|---|---|---|---|
| GCA_963645785.1 | o__RFP12;f__UBA1067 | 1790427 | 1707 | 76.21 |
| GCA_963659085.1 | o__RFP12;f__UBA1067 | 1162374 | 1194 | 71.29 |
| GCA_963661055.1 | o__RFP12;f__UBA1067 | 1882898 | 1619 | 91 |
| GCA_002309765.1 | o__RFP12;f__UBA1067 | 3673306 | 2628 | 86.05 |
| GCA_017454745.1 | o__RFP12;f__UBA1067 | 3858549 | 2705 | 88.64 |
| GCA_002309585.1 | o__RFP12;f__UBA1067 | 3781978 | 2828 | 57.66 |
| GCA_002317575.1 | o__RFP12;f__UBA1067 | 2795861 | 2235 | 70.12 |
| GCA_017521065.1 | o__RFP12;f__UBA1067 | 3520818 | 3263 | 53.11 |
| GCA_017936245.1 | o__RFP12;f__UBA1067 | 3541139 | 3177 | 56.09 |
| GCA_024405335.1 | o__RFP12;f__UBA1067 | 2070756 | 1775 | 80.11 |
| GCA_902779565.1 | o__RFP12;f__UBA1067 | 4093499 | 3198 | 74.82 |
| GCA_934667425.1 | o__RFP12;f__UBA1067 | 3272908 | 2441 | 84.97 |
| GCA_946631345.1 | o__RFP12;f__UBA1067 | 3024164 | 2515 | 55 |
| GCA_963618095.1 | o__RFP12;f__UBA1067 | 1690698 | 1598 | 69.8 |
| GCA_963626295.1 | o__RFP12;f__UBA1067 | 1966951 | 1829 | 82.56 |
| GCA_963653155.1 | o__RFP12;f__UBA1067 | 3215061 | 2563 | 85.49 |
| GCA_009777165.1 | o__RFP12;f__UBA1067 | 2450500 | 2115 | 65.41 |
| GCA_009778475.1 | o__RFP12;f__UBA1067 | 3966866 | 2852 | 79.08 |
| GCA_949285835.1 | o__RFP12;f__UBA3636 | 2671747 | 2160 | 91.13 |
| GCA_012514445.1 | o__RFP12;f__UBA3636 | 3266551 | 3001 | 64.7 |
| GCA_012511845.1 | o__RFP12;f__UBA3636 | 2623161 | 2487 | 77.21 |
| GCA_022837245.1 | o__RFP12;f__UBA3636 | 3158365 | 2925 | 62.14 |
| GCA_016295275.1 | o__RFP12;f__UBA3636 | 2340517 | 1906 | 75.1 |
| GCA_949295265.1 | o__RFP12;f__UBA3636 | 2823886 | 2178 | 89.09 |
| GCA_038325415.1 | o__RFP12;f__UBA3636 | 1996229 | 1924 | 58.2 |
| GCA_017424105.1 | o__RFP12;f__UBA3636 | 1778250 | 1441 | 78.43 |
| GCA_017481725.1 | o__RFP12;f__UBA3636 | 2593024 | 2007 | 85.4 |
| GCA_017648065.1 | o__RFP12;f__UBA3636 | 2060779 | 1630 | 84.62 |
| GCA_937911295.1 | o__RFP12;f__UBA3636 | 2596783 | 2063 | 89.99 |
| GCA_002448475.1 | o__RFP12;f__UBA3636 | 4477447 | 3293 | 89.38 |
| GCA_017413145.1 | o__RFP12;f__UBA3636 | 3286361 | 2889 | 69.8 |
| GCA_017450425.1 | o__RFP12;f__UBA3636 | 1358022 | 1363 | 52.86 |
| GCA_017457625.1 | o__RFP12;f__UBA3636 | 2566374 | 2165 | 75.05 |
| GCA_017476965.1 | o__RFP12;f__UBA3636 | 2088857 | 1989 | 72.51 |
| GCA_902761575.1 | o__RFP12;f__UBA3636 | 3405367 | 2868 | 91.95 |
| GCA_902793555.1 | o__RFP12;f__UBA3636 | 3379398 | 2699 | 79.91 |
| GCA_946534105.1 | o__RFP12;f__UBA3636 | 3931963 | 3046 | 71.23 |

Supplemental Table 1: Metadata and taxonomy of species representative Kiritimatiellia genomes, Continued

| GTDB Accession | GTDB Taxonomy | Genome Size | # CDS | Q-Score |
|---|---|---|---|---|
| GCA_946534745.1 | o__RFP12;f__UBA3636 | 2079032 | 1851 | 65.67 |
| GCA_946612175.1 | o__RFP12;f__UBA3636 | 2993096 | 2442 | 82.39 |
| GCA_946624575.1 | o__RFP12;f__UBA3636 | 3947021 | 3089 | 91.13 |
| GCA_947169825.1 | o__RFP12;f__UBA3636 | 3081420 | 2486 | 67.59 |
| GCA_028708815.1 | o__RZYC01;f__RZYC01 | 4023004 | 3244 | 90.64 |
| GCA_009930475.1 | o__RZYC01;f__RZYC01 | 4773275 | 3997 | 83.51 |
| GCA_009929845.1 | o__RZYC01;f__RZYC01 | 5644885 | 4645 | 79.4 |
| GCA_018814445.1 | o__SLAD01;f__JAHIZV01 | 4062274 | 3450 | 89.16 |
| GCA_007127595.1 | o__SLAD01;f__SLAD01 | 2325650 | 2263 | 65.46 |
| GCA_007128275.1 | o__SLAD01;f__SLAD01 | 2799226 | 2558 | 83.55 |
| GCA_007136425.1 | o__SLAD01;f__SLAD01 | 2601423 | 2320 | 86.45 |
| GCA_007123265.1 | o__SLAD01;f__SLAD01 | 2430054 | 2229 | 88.12 |
| GCA_007125875.1 | o__SLAD01;f__SLAD01 | 2372393 | 2176 | 85.22 |
| GCA_007126515.1 | o__SLAD01;f__SLAD01 | 2465461 | 2318 | 88.49 |
| GCA_007126995.1 | o__SLAD01;f__SLAD01 | 2466917 | 2371 | 66.9 |
| GCA_007134735.1 | o__SLAD01;f__SLAD01 | 2584683 | 2209 | 81.14 |
| GCA_003562335.1 | o__SLAD01;f__SLAD01 | 2659874 | 2510 | 91.68 |
| GCA_007134175.1 | o__SLAD01;f__SLAD01 | 2352921 | 2268 | 75.95 |
| GCA_007130845.1 | o__SLAD01;f__SLAD01 | 2218741 | 2163 | 80.71 |
| GCA_012103575.1 | o__SLAD01;f__WLWN01 | 3136281 | 3622 | 91.5 |
| GCA_963578625.1 | o__SLOV01;f__SLOV01 | 5314967 | 4771 | 73.65 |
| GCA_007132345.1 | o__SLOV01;f__SLOV01 | 3526840 | 3180 | 86.94 |
| GCA_037134535.1 | o__SPBP01;f__SPBP01 | 3725035 | 3564 | 79.66 |
| GCA_037135275.1 | o__SPBP01;f__SPBP01 | 3702885 | 3367 | 69.22 |
| GCA_032123355.1 | o__SPBP01;f__SPBP01 | 3350828 | 3178 | 89.65 |
| GCA_003566475.1 | o__SS1-B-03-39;f__PXDE01 | 4419250 | 4070 | 75.39 |
| GCA_029936175.1 | o__SS1-B-03-39;f__UBA6053 | 5040440 | 4213 | 79.36 |
| GCA_003565035.1 | o__SS1-B-03-39;f__UBA6053 | 3885734 | 3522 | 80.63 |
| GCA_003565815.1 | o__SS1-B-03-39;f__UBA6053 | 3655480 | 3344 | 77.98 |
| GCA_020831625.1 | o__SS1-B-03-39;f__UBA6053 | 4885981 | 3874 | 90.44 |
| GCA_020831635.1 | o__SS1-B-03-39;f__UBA6053 | 6008866 | 4878 | 96.1 |
| GCA_002429065.1 | o__SS1-B-03-39;f__UBA6053 | 5686037 | 4737 | 94.72 |
| GCA_018401235.1 | o__SS1-B-03-39;f__UBA6053 | 2492457 | 2453 | 82.79 |
| GCA_029269125.1 | o__SS1-B-03-39;f__UBA6053 | 4524781 | 3860 | 92.38 |
| GCA_029719485.1 | o__SS1-B-03-39;f__UBA6053 | 4432027 | 3731 | 89.63 |
| GCA_040905625.1 | o__SS1-B-03-39;f__UBA6053 | 2993956 | 2703 | 66.9 |
| GCA_937127435.1 | o__SS1-B-03-39;f__UBA6053 | 5205169 | 4590 | 82.76 |

Supplemental Table 1: Metadata and taxonomy of species representative Kiritimatiellia genomes, Continued

| GTDB Accession | GTDB Taxonomy | Genome Size | # CDS | Q-Score |
|---|---|---|---|---|
| GCA_002429005.1 | o__SS1-B-03-39;f__UBA6053 | 5778707 | 4880 | 89.52 |
| GCA_937920455.1 | o__UBA8416;f__CALETX01 | 2297498 | 2204 | 60.59 |
| GCA_945883775.1 | o__UBA8416;f__CAMCYG01 | 2657872 | 2453 | 88.2 |
| GCA_030632465.1 | o__UBA8416;f__CAMZLY01 | 1973045 | 1869 | 69.74 |
| GCA_947311825.1 | o__UBA8416;f__CAMZLY01 | 2497075 | 2199 | 87.32 |
| GCA_013203785.1 | o__UBA8416;f__JABMPJ01 | 3995952 | 3462 | 88.98 |
| GCA_016783675.1 | o__UBA8416;f__JABMPJ01 | 2615471 | 2258 | 66.15 |
| GCA_018648805.1 | o__UBA8416;f__JABMPJ01 | 4848971 | 3791 | 80.49 |
| GCA_029939545.1 | o__UBA8416;f__JABMPJ01 | 2599436 | 2424 | 86.66 |
| GCA_030745755.1 | o__UBA8416;f__JABMPJ01 | 3068618 | 2575 | 87.72 |
| GCA_016783565.1 | o__UBA8416;f__JADHWI01 | 2418961 | 2146 | 84.38 |
| GCA_032123415.1 | o__UBA8416;f__JADHWI01 | 2879433 | 2985 | 61.37 |
| GCA_030744755.1 | o__UBA8416;f__JADHWI01 | 6279599 | 4981 | 76.93 |
| GCA_016929365.1 | o__UBA8416;f__JAFGIZ01 | 2379530 | 2163 | 69.76 |
| GCA_016934225.1 | o__UBA8416;f__JAFGTH01 | 2131810 | 2058 | 71.76 |
| GCA_021413965.1 | o__UBA8416;f__JAIOPI01 | 5430970 | 4174 | 90.03 |
| GCA_027620105.1 | o__UBA8416;f__JAIOPI01 | 2385131 | 2452 | 57.99 |
| GCA_945949405.1 | o__UBA8416;f__JAIOPI01 | 1955417 | 1661 | 75.66 |
| GCA_028822745.1 | o__UBA8416;f__JAPKCZ01 | 3377702 | 3410 | 62.74 |
| GCA_032123435.1 | o__UBA8416;f__JAQBFJ01 | 2574334 | 2484 | 70.83 |
| GCA_041306605.1 | o__UBA8416;f__JAQBFJ01 | 2937975 | 2735 | 83.31 |
| GCA_030590485.1 | o__UBA8416;f__JAUUYL01 | 2625645 | 2467 | 87.51 |
| GCA_041602695.1 | o__UBA8416;f__JBFIFS01 | 3002913 | 2809 | 77.43 |
| GCA_003565095.1 | o__UBA8416;f__PXDI01 | 3227609 | 2859 | 86.42 |
| GCA_030018915.1 | o__UBA8416;f__SLMC01 | 2118169 | 1918 | 88.95 |
| GCA_007133745.1 | o__UBA8416;f__SLMC01 | 2620838 | 2304 | 77.99 |
| GCA_007134465.1 | o__UBA8416;f__SLMC01 | 3987713 | 3391 | 74.5 |
| GCA_018645275.1 | o__UBA8416;f__UBA8416 | 4685755 | 4201 | 77.99 |
| GCA_003567005.1 | o__UBA8416;f__UBA8416 | 3701402 | 3093 | 75.75 |
| GCA_007126175.1 | o__UBA8416;f__UBA8416 | 1328710 | 1271 | 66.24 |
| GCA_007129405.1 | o__UBA8416;f__UBA8416 | 1968878 | 1834 | 71.19 |
| GCA_002428045.1 | o__UBA8416;f__UBA8416 | 4227307 | 3735 | 70.67 |
| GCA_020048935.1 | o__UBA8416;f__UBA8416 | 2393843 | 2157 | 67.86 |
| GCA_016873655.1 | o__UBA8416;f__VGWI01 | 3075852 | 2781 | 86.1 |
| GCA_016873635.1 | o__UBA8416;f__VGWJ01 | 3237298 | 2768 | 90.06 |
| GCA_001804885.1 | o__UBA8416;f__YA12-FULL-48-11 | 5678892 | 4926 | 76.7 |
| GCA_028716365.1 | o__UBA8416;f__YA12-FULL-48-11 | 5799834 | 4515 | 85.6 |
| GCA_001803315.1 | o__UBA8416;f__YA12-FULL-60-10 | 4090690 | 3418 | 83.7 |
